# Simulation-guided non-thermal low-intensity ultrasound reprograms the tumor immune microenvironment and engages systemic antitumor immunity in a syngeneic orthotopic mouse model of breast cancer

**DOI:** 10.64898/2026.08.13.743931

**Authors:** Reyhaneh Hooshmandabbasi, Ali Kazemian, Rahul Singha, Manuel Vielma Blanco, Niloufar Nikkhah Bahrami, Thomas Hauser, Mathias S. Weyland, Franco Guscetti, David Wahl, Daniel Fehr, Mathias Bonmarin, Stephan Scheidegger, Caroline Maake

**Author notes:** **Corresponding author:** Reyhaneh Hooshmandabbasi, DVM, PhD.

## Abstract

**Introduction:** Therapeutic ultrasound has been extensively studied in ablative and sonodynamic contexts, leaving the intrinsic bioactivity of continuous non-thermal low-intensity ultrasound (LIU) largely uncharacterized.

**Objectives:** To characterize the tumor biological and immunomodulatory effects of non-thermal continuous LIU in complementary *in vitro* and *in vivo* breast cancer models, underpinned by a standardized exposure platform characterized through finite element simulations and experimental validation.

**Methods:** Acoustic and thermal fields were characterized and optimized using *in silico* simulations and validated against hydrophone and temperature measurements to ensure homogeneous, non-thermal exposure (1MHz, 1W/cm^2^, 100% duty cycle). 4T07 murine mammary carcinoma spheroids received 20min LIU treatment, and metabolic activity, apoptosis, and intracellular stress-associated markers were assessed. In a syngeneic orthotopic 4T07 mammary carcinoma model in BALB/c mice, up to six LIU treatment cycles were administered; tumor growth, survival, histopathology, immunohistochemistry, bulk tumor RNA sequencing, spleen volume and plasma cytokine profiles were assessed.

**Results:** *In vitro* and intratumoral temperatures remained within the physiological range (≤39°C) throughout exposure. In spheroids, LIU reduced ATP content by more than 40% and significantly increased apoptotic, Hsp70⁺ and Hsp90⁺ cell fractions. *In vivo*, cyclic LIU slowed tumor growth, increased intratumoral necrosis, and significantly prolonged time to humane endpoint compared to untreated controls. LIU promoted early intratumoral myeloid cell infiltration and shifted the tumor transcriptome (2,573 differentially expressed genes), with enrichment in gene sets associated with immunogenic cell death, pattern-recognition, inflammatory, and innate and adaptive immune programs and downregulation of pro-tumorigenic pathways. LIU enriched the transcriptional signatures of M1 macrophage polarization and, notably, B-cell compartment engagement, which has not previously been reported for standalone continuous mechanical ultrasound. LIU significantly attenuated tumor-associated splenomegaly and elevated plasma IL-1α, TNF-α, and IL-10.

**Conclusion:** These results establish a reproducible preclinical platform and provide a hypothesis-generating mechanistic basis for evaluating LIU as an adjunct to immune checkpoint blockade.

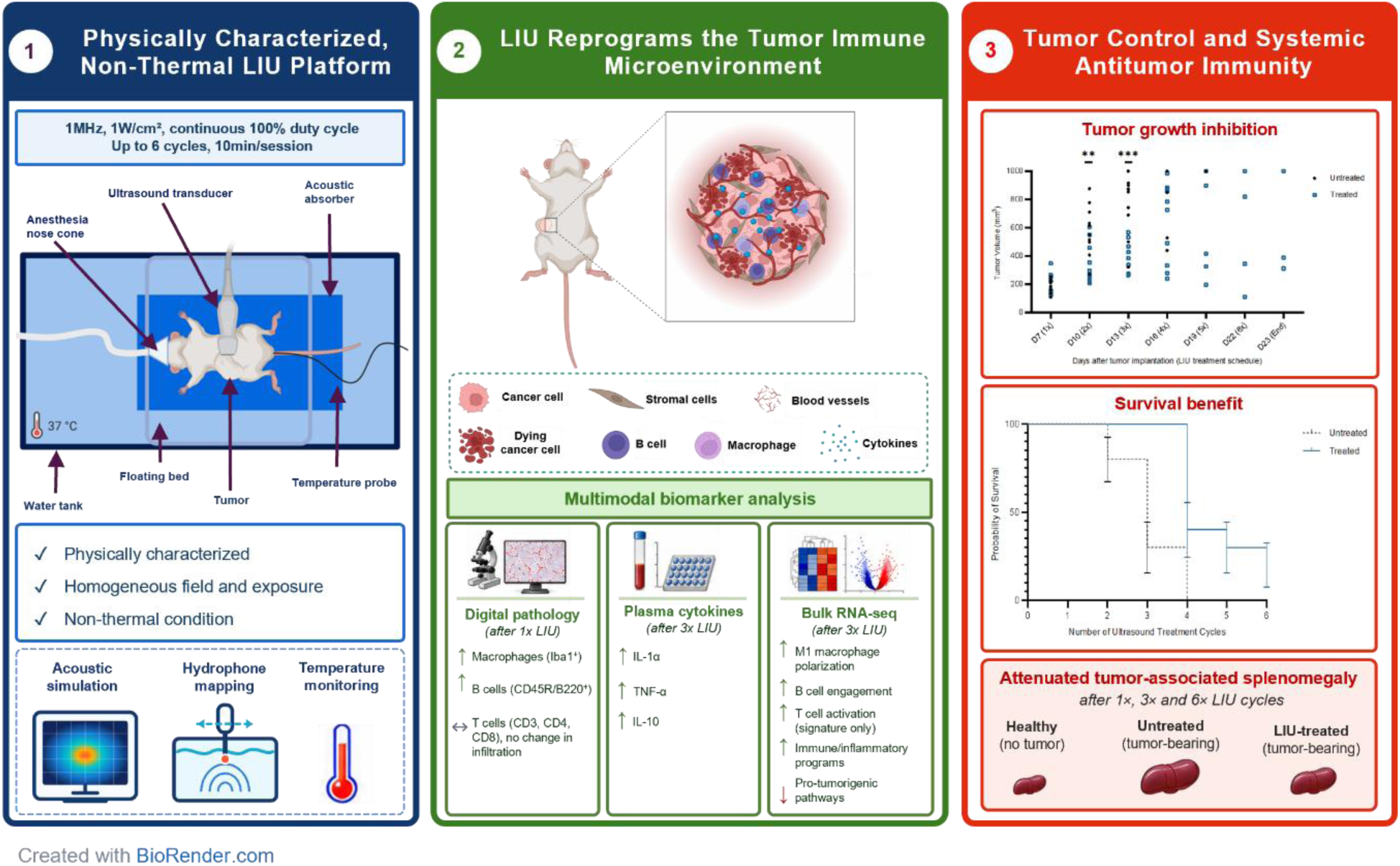

## Introduction

Breast cancer is the most frequently diagnosed malignancy in women and a leading cause of cancer-related death worldwide [1]. Despite continued progress in surgery, radiotherapy, chemotherapy and endocrine therapy, durable control of recurrent, locally advanced and metastatic disease remains difficult to achieve, because tumors acquire treatment resistance and actively evade immune surveillance [2,3]. Immune checkpoint blockade has transformed the management of several malignancies [4], yet in breast cancer its benefit is restricted to a minority of patients, a limitation widely attributed to the predominantly immunologically “cold”, immune-excluded and immunosuppressive nature of the breast tumor microenvironment (TME) [5]. Reprogramming such cold tumors into immunologically active, “hot” lesions, without adding the systemic toxicity of further pharmacological agents, has consequently become a central objective of modern oncology.

Locally delivered, energy-based physical stimuli may offer one route to this conversion: by damaging tumor cells *in situ*, exposing tumor antigens and releasing endogenous danger signals, they can inflame and remodel the TME without the off-target burden of systemic therapy [6]. Among such modalities, ultrasound is especially attractive because it is non-ionizing, non-invasive, widely available and, uniquely among physical energy sources, capable of transmitting mechanical energy through intact tissue and concentrating it at deep-seated targets with millimetric precision [7]. While high-intensity focused ultrasound (HIFU) ablates tumors through thermal coagulative necrosis and is already established in clinical practice [8], interest has increasingly shifted toward low-intensity ultrasound (LIU; typically 0.1-3 W/cm^2^), which acts predominantly through non-thermal mechanical bioeffects [9]. Phenomena such as stable cavitation, acoustic microstreaming and oscillatory radiation forces impose mechanical stresses on cells, engaging mechanotransduction pathways and eliciting responses including altered membrane permeability, cytoskeletal remodeling and changes in signal transduction. Critically, focused LIU has been reported to induce surface translocation of Hsp70 and Hsp90 and release of HMGB1 in breast and prostate cancer cells, implicating immunogenic signaling through damage-associated molecular pattern (DAMP) release [10]. These observations position non-thermal ultrasound as a potential mechano-immunological stimulus capable of eliciting strong immune responses, a concept increasingly explored under the umbrella of ultrasound-assisted immunotherapy [11].

Despite this promise, the evidence base for continuous LIU in solid tumors remains limited. Existing studies have largely employed LIU as an adjunct to drug delivery or mild hyperthermia, or have been confined to *in vitro* settings or combined with thermal or pharmacological agents [12–14]. Consequently, the intrinsic, drug-free biological activity of continuous LIU, and above all its capacity to modulate antitumor immunity *in vivo*, has never been systematically and mechanistically characterized under rigorously controlled non-thermal conditions.

A compounding obstacle is the lack of standardized acoustic exposure protocols across the field. Intensities, frequencies and sonication geometries are reported inconsistently, and treatment parameters are frequently established empirically, introducing uncontrolled confounders such as unintended heating or heterogeneous cavitation activity [15–17]. Continuous LIU is uniquely suited to address this: it delivers a sustained, spatially uniform mechanical stimulus that can be standardized and reproduced without cavitation control agents or intermittent pulsing. To achieve this, we combined acoustic and thermal *in silico* simulations with experimental validation to establish a platform delivering homogeneous, non-thermal and reproducible exposure in both *in vitro* and *in vivo* settings. For *in vivo* characterization, we selected the murine 4T07 syngeneic orthotopic breast cancer model, in which an intact immune system is required to restrain tumor growth [18], providing a physiologically relevant testbed for immune-activating intervention.

Building on this platform, the present study provides a mechanistically defined characterization of the non-thermal, mechanically driven biological and immunomodulatory effects of continuous LIU in breast cancer. We asked whether a rigorously controlled, strictly non-thermal LIU exposure can induce immunogenic stress responses, remodel tumor architecture and reprogram the TME toward an antitumor immune state, and whether these local effects are associated with systemic immune responses that may contribute to antitumor immunity and improved tumor control.

## Materials and methods

If not otherwise mentioned, all chemicals were obtained from Sigma Aldrich, Buchs, Switzerland.

### Cell culture

4T07 cells [19] were cultured in Dulbecco’s Modified Eagle’s Medium (ATCC, Manassas, USA), penicillin/streptomycin (50U/ml each, ThermoFisher Scientific, Waltham, USA) and 5% fetal calf serum (FCS, ThermoFisher Scientific) at 37°C in a humidified atmosphere with 5% CO_2_. Cells tested negative for mycoplasma and mouse pathogens (IMPACT II Test, IDEXX Bioanalytics, Westbrook, USA) and were used in passages 3-4.

4T07 spheroids were generated using 3D CoSeedis™ Chip200 (abc biopply, Solothurn, Switzerland) according to the manufacturer’s protocol. Briefly, 2000 cells were seeded per microwell to generate spheroids with 300-500µm diameter. Spheroids were imaged with GelCount^TM^ (Oxford Optronix, Abingdon, UK) and analyzed with ImageJ 1.48v (National Institutes of Health, Bethesda, USA). The spheroid volume V was calculated with the formula V= 4π/3(√(A/π))^3^.

### *In silico* determination of *in vitro* ultrasound conditions

Acoustic pressure distribution within the microplate and the optimal transducer-to-sample distance were determined at 1MHz and 100% duty cycle using the k-Wave MATLAB toolbox for time-domain simulation of acoustic wave fields [20,21]. Simulations were run on the University of Zurich ScienceCluster high-performance computing system. The microplate, water tank, and transducer geometries were modelled using Autodesk Fusion 360 (Autodesk, Inc., San Francisco, CA, USA) (**Fig. S1**).

### *In vitro* ultrasound setup

The custom *in vitro* ultrasound setup (**Fig. 1; S1.1**) consisted of a water tank, microplate holder, acoustic absorber, heater, and customized ultrasound system. The ultrasound device consisted of a transducer (SMMSG25F1000, STEMiNC, Miami, USA) featuring a stainless-steel housing with a diameter of approx. 40mm. The transducer’s piezo element, a disk made of modified PZT-4 material with a diameter of 25mm mounted to the stainless-steel housing, had a nominal resonance frequency of 1MHz. An Aptflex F36 acoustic absorber positioned above the microplates reduced reflections from the water–air interface and minimized standing-wave formation [22]. Details of the setup fabrication, control electronics, operating parameters, and output-linearity assessment are provided in the Supplementary Methods (**S1.1 and S1.2**).

**Fig. 1.**
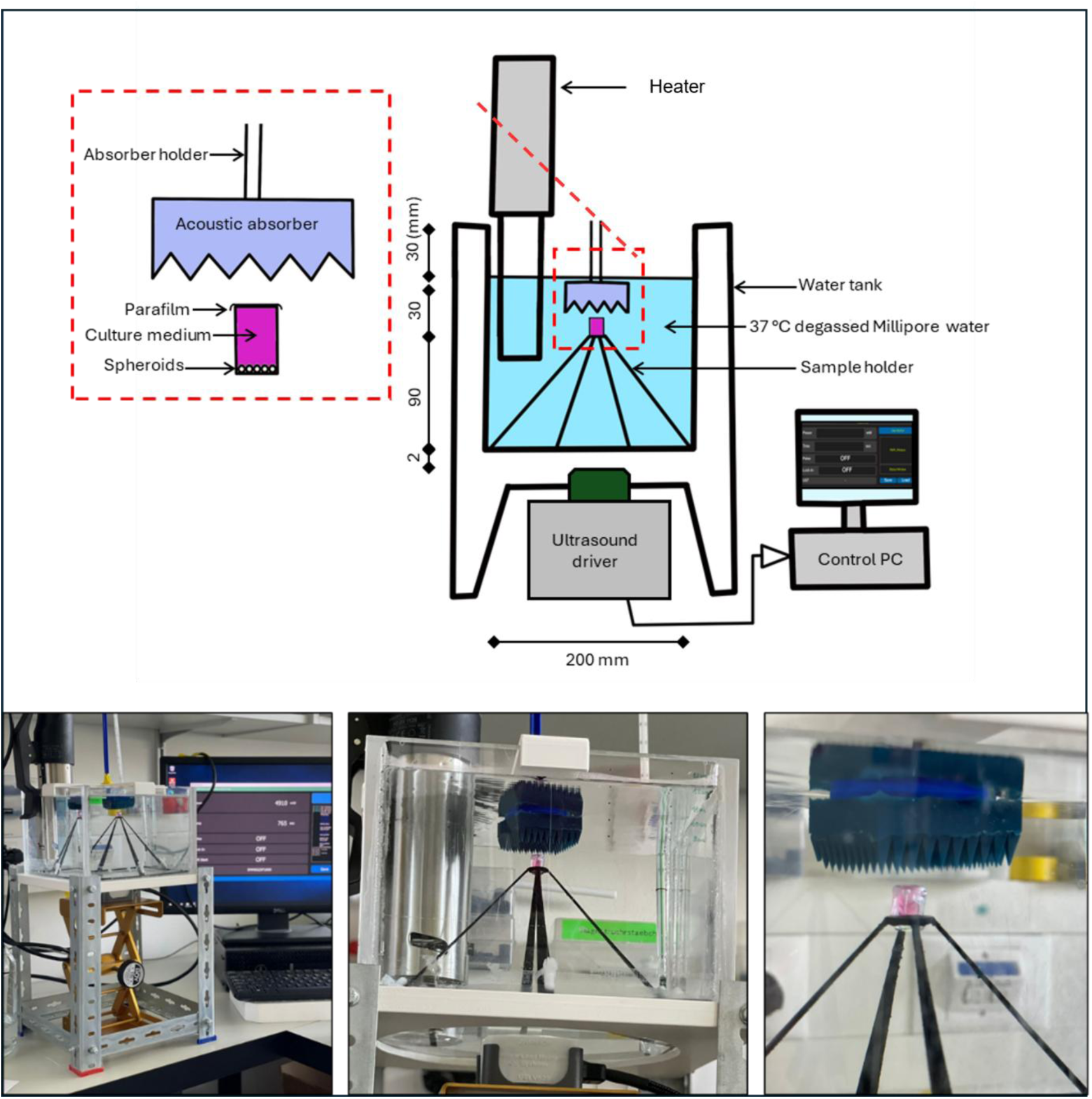
Experimental setup for in vitro low-intensity ultrasound (LIU) experiments. 4T07 spheroids (x8 per microplate) were placed in sealed microplates and positioned inside a water tank containing degassed MilliQ water at 37°C. Ultrasound was delivered using a customized 1MHz transducer driven by a control PC, with acoustic absorption provided by an absorber plate to minimize reflections. LIU conditions: 1MHz, 1W/cm^2^, 100% duty cycle, 20min.

Hydrophone measurements were performed at 94mm from the tank bottom, corresponding to the bottom of the microplate, using a needle hydrophone (Müller-Platte Nadelsonde, Müller Instruments, Oberursel, Germany). The temperature was monitored using a thermocouple sensor (IT-18 Flexible Microprobe, Physitemp Instruments, Clifton, USA) positioned at the center of the microplate. All measurements were performed under identical acoustic conditions (1MHz, 1W/cm^2^, 100% duty cycle, 20min.). In the following, 100% duty cycle refers to a continuous exposure during the indicated treatment time.

### *In vitro* ultrasound treatment

Spheroids (300-500µm; 8 per condition, pooled per experiment) were transferred to a flat bottom Corning Stripwell microplate (diameter: 6.9mm; Corning, NY, USA). The microplate was completely filled with cell culture medium, sealed with Parafilm (Electron Microscopy Sciences, Hatfield, PA, USA) and placed in the sample holder within the water tank (94mm from tank base to the microplate bottom). Spheroids were sonicated once (1MHz, 1W/cm^2^, 100% duty cycle, 20min), followed by incubation under cell culture conditions with fresh medium for 24h. Morphological characteristics of spheroids were assessed by an Olympus CKX53 inverted microscope (Olympus, Tokyo, Japan). Untreated spheroids (n=8) served as controls.

### CellTiter-Glo assay

Metabolic activity of spheroids 24h after ultrasound treatment was analyzed with the CellTiter-Glo® 3D Viability Assay (Promega, Madison, WI, USA) as recommended by the manufacturer. Briefly, after ultrasound experiments as above, medium was replaced by 200µL/well CellTiter-Glo® reagent. Luminescence was measured after 12min using an Agilent BioTek Cytation 5 reader (Agilent Technologies, Santa Clara, CA, USA).

### Flow cytometry

Ultrasound-treated and untreated spheroids (8 per condition, pooled per experiment) were washed and centrifuged (5min, 350×g). Pellets were resuspended in 200µL Accumax (ThermoFisher Scientific) and transferred each to a mixture of 400µL Accumax and 600µL collagenase IV (1mg/mL; Merck, Darmstadt, Germany). The suspensions were incubated at 37°C for 20min, washed with PBS and filtered through a 70μm cell strainer (BD Biosciences). Samples were transferred to V-bottom 96-well plates (ThermoFisher Scientific) and stained with Zombie UV™ viability dye (1:300, BioLegend, San Diego, USA) for 20min at 4°C in the dark, followed by APC/Fire™ 750-conjugated Annexin V (1:10, BioLegend) and Alexa Fluor® 594-conjugated anti-Calreticulin antibody (D3E6, 1:100, Cell Signaling Technology, Danvers, USA) for another 20min at 4°C in the dark. Subsequently, cells were fixed and permeabilized using the fixation/permeabilization solution from the Cytofix/Cytoperm™ kit (BD Biosciences) according to the manufacturer’s instructions. Intracellular staining was performed for 30min at 4°C in the dark using an antibodies cocktail in Perm/Wash buffer containing PE-conjugated Hsp70/Hsp72 (C92F3A, 1:100, Enzo Life Sciences, Farmingdale, USA), DyLight™ 488-conjugated Hsp90 (AC88, 1:100, Enzo Life Sciences), and Alexa Fluor® 647-conjugated HMGB1 (3E8, 1:100, BioLegend). Cells were washed and resuspended in 200µL Perm/Wash buffer for data acquisition in a spectral flow cytometer (Cytek Aurora 5L, Cytek Biosciences, Fremont, USA). SpectroFlo® software (Cytek Biosciences) was used for unmixing. Data analysis, including gating strategies, was conducted with FlowJo™ v10.9.4 (BD Biosciences). The proportion of total cells positive for Hsp70, Hsp90, HMGB1, and calreticulin was quantified independent of Annexin V/Zombie UV status. Unstained and single-stained samples served as controls.

### Pre-clinical model

Female BALB/c wild-type mice (n=72, 12 weeks old, Charles River, Sulzfeld, Germany) were housed under pathogen-free conditions at the Laboratory Animal Services Center at the University of Zurich, had *ad libitum* access to food and water and were maintained on a 12h light/dark cycle with environmental enrichment.

Six mice were assigned to a control group, while 66 mice received a 50µL injection of 100,000 4T07 cells into the right fourth mammary fat pad. Tumor growth was measured every third day using caliper. Tumor volume (mm^3^) was calculated as: (d^2^ × D)/2, (d = tumor width; D = tumor length). Mice were euthanized by CO₂ inhalation.

### Ethics statement

All animal procedures were conducted in accordance with the ethical policies and protocols approved by the Cantonal Veterinary Office Zurich (ZH 062/2022).

### *In silico* determination of *in vivo* ultrasound conditions

Autodesk Fusion 360 (Autodesk Inc.) and published segmentation data were used to construct a 3D mouse model [23]. The experimental setup and the location of the tumor was also modeled in Autodesk Fusion 360 (**Fig. S2**). Physical properties of the different layers were extracted from the tissue database of the IT’IS Foundation, Zurich, Switzerland [24].

### *In vivo* ultrasound setup

A schematic picture of the setup is provided in **Fig. 2**. It consisted of a plexiglass animal holder placed in a water bath (water degassed, 37°C) equipped with an Aptflex F28 ultrasound absorber (Precision Acoustics, Dorchester, UK) at the tank bottom. The transducer of an EMS Primo Therasonic 460 device (EMS Physio) was held in place by a robotic arm (RS PRO, Beauvais, France) throughout the treatment. Based on *in silico* calculations (see above), a 20mm-thick gel pad (Aquaflex, Parker Laboratories, Fairfield, USA) was positioned between the transducer and the tumor.

**Fig. 2.**
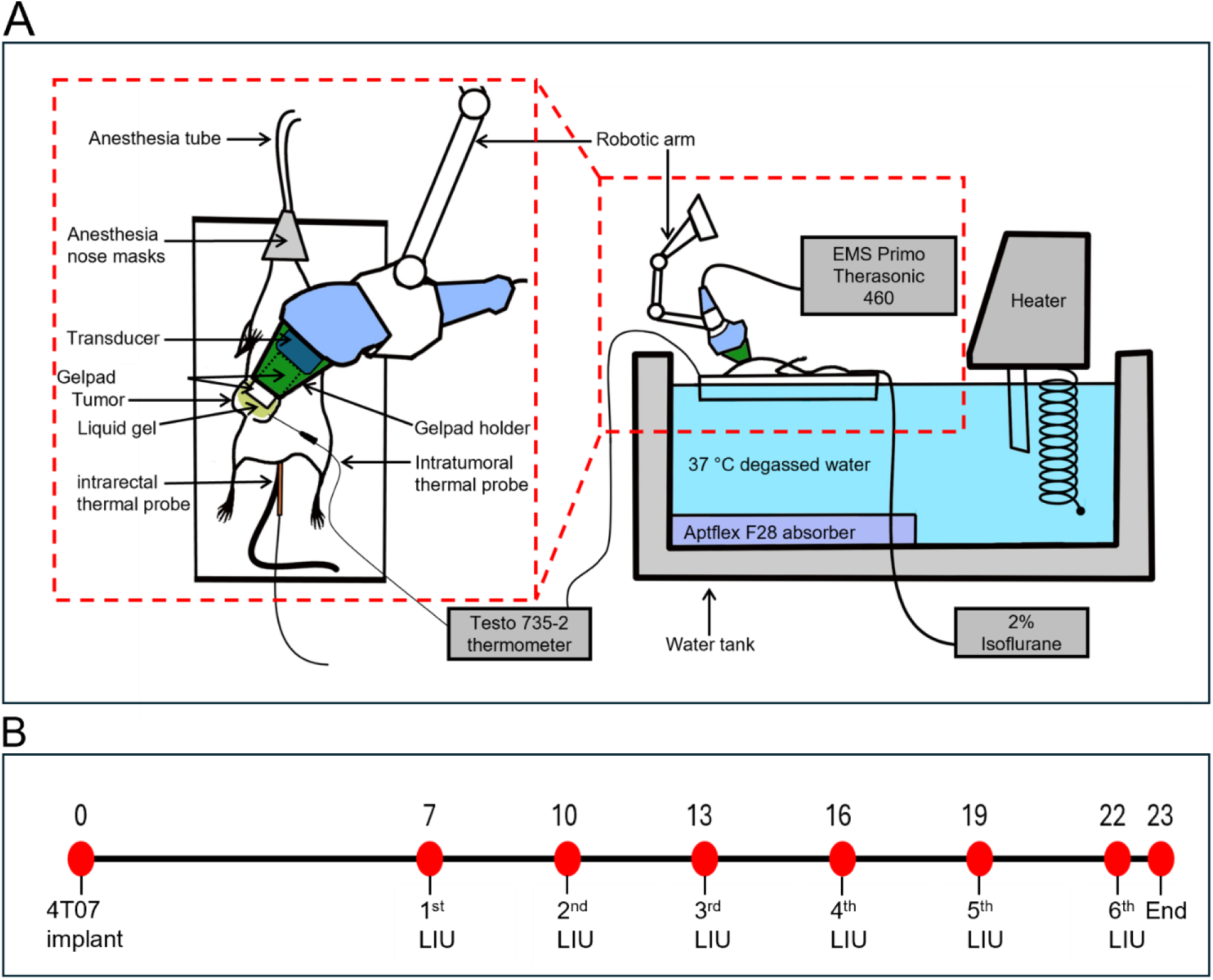
In vivo low-intensity ultrasound (LIU) experimental setup and treatment scheme. (A) Schematic of the customized setup used for ultrasound delivery to orthotopic mammary tumors in mice. Animals were anesthetized with isoflurane and placed supine on the sample holder, positioned above the water tank containing degassed water at 37°C. A 1MHz transducer was coupled to the tumor surface through a 20mm gel pad, with an absorber included to minimize acoustic reflections. Temperature of the body and center of the tumor was continuously monitored via rectal and intratumoral probes, respectively. (B) Experimental timeline of LIU treatment. 4T07 cancer cells were implanted on Day 0. LIU was applied (1MHz, 1W/cm^2^, 100% duty cycle, 10min.) starting on Day 7 and repeated every third day for up to six sessions (Days 7, 10, 13, 16, 19, and 22). Remaining animals were euthanized 24h after the last treatment (Day 23). Histopathology was performed after the 1st LIU, and RNA sequencing and plasma cytokine profiling after the 3rd LIU.

### *In vivo* ultrasound treatment

Upon reaching a tumor volume of at least 100mm^3^, mice were stratified by tumor size and randomized into untreated (n=30) or ultrasound therapy (n=36) groups. After isoflurane anesthesia (5% for induction, 2% for maintenance), mice were shaved, positioned supine on the animal holder and the tumor site was coated with Aquasonic 100 Ultrasonic Gel (Parker Laboratories). Ultrasound treatments (1MHz, 1W/cm^2^, 100% duty cycle, 10min.) were repeated up to 6 cycles under isoflurane anesthesia at 72h intervals. Tumor temperature was recorded using a MT-29/5HT needle microprobe sensor (Physitemp, Clifton, USA) inserted into the tumor center, connected to a Testo 735-2 thermometer (Testo AG, Mönchaltorf, Switzerland). Rectal temperature was monitored using the PhysioSuite system (Kent Scientific, Torrington, USA). Untreated mice underwent tumor caliper measurements on the same schedule as the treatment group but did not receive isoflurane anesthesia or ultrasound exposure.

### Histopathological and immunohistochemical analyses

Following euthanasia, tumors collected 24h after the first ultrasound cycle and untreated tumors at the same time point were fixed in 10% neutral-buffered formalin for 24h, embedded in paraffin and sectioned (2.5µm).

For histopathology, sections were stained with hematoxylin and eosin (H&E) according to standard protocols. For immunohistochemistry, sections were treated with Peroxidase Blocking Reagent (S2023, Agilent Technologies) for 10min at room temperature (RT), followed by antigen retrieval for 20min using the PT Link system (Agilent Technologies) with citrate buffer (pH 6.0, Agilent Technologies) for Iba1, CD45R and CD8, and with EDTA buffer (pH 9.0, Agilent Technologies) for CD3 and CD4.

Sections were incubated with rabbit anti-Iba1 (1:1000, 019-19741, FUJIFILM Wako Pure Chemical Corporation, Osaka, Japan; 1h at RT), rabbit anti-CD3 (1:250, ab16669, Abcam, Cambridge, UK; overnight at 4°C), rabbit anti-CD4 (1:500, ab183685, Abcam; 1h at RT), rabbit anti-CD8 (1:400, 98941, Cell Signaling Technology; overnight at 4°C), or rat anti-CD45R (1:800, 553084, BD Pharmingen; 1h at RT). For sections stained with rat anti-CD45R, a rabbit anti-rat bridging antibody (Vector BA-4000; 1:1000, 30min at RT) was applied. Thereafter, slides were treated for 30min at RT with the EnVision+ System-HRP for rabbit (K4003, Agilent Technologies) followed by the AEC single solution (Zytomed, ZUC037-100) for 10min and counterstaining with hematoxylin for 2sec.

Following whole-slide scanning (NanoZoomer 2.0-HT, Hamamatsu, Hamamatsu city, Japan), image analysis was conducted using Visiopharm software (Visiopharm, Broomfield, CO, USA). Tumor destruction rate was calculated as the proportion of necrotic tissue relative to the total tumor area in H&E-stained sections. In immunohistochemically stained sections, AEC-positive areas were quantified and normalized to the total tumor area. Per tumor, 3-5 sections were evaluated and averaged to yield a single value per animal, which was plotted as one data point per biological replicate in the corresponding figures. All analyses were conducted blinded to experimental groups.

### Multiplex cytokine analyses

Blood was collected from healthy control, untreated tumor-bearing, and treated tumor-bearing mice (n=3 per group) 24h after the third treatment cycle into BD Vacutainer® EDTA Tubes (Becton Dickinson, Franklin Lakes, NJ, USA) and centrifuged (1,600xg, 15min at RT and 16,000xg, 10min at 4°C). Cytokine concentrations in plasma were quantified using the Mouse Inflammation Panel (13-plex, V-bottom plate; BioLegend) following the manufacturer’s instructions. TNF-α, IL-1α, IL-1β, IL-12p70, IL-10, IL-6, IL-17A, IL-23, IL-27, IFN-β, IFN-γ, MCP-1, and GM-CSF were measured in triplicate using a Cytek Aurora flow cytometer (Cytek Biosciences) with phycoerythrin (PE) as the reporter and allophycocyanin (APC) as the bead channel. Data were analyzed using the LEGENDplex Data Analysis Software (BioLegend).

### RNA sequencing

Tumors (200–500mg) were collected from mice (n=3 per group) 24h after the third treatment cycle, preserved in RNAprotect (Qiagen, Hilden, Germany) and submitted to Qiagen Genomic Services. Briefly, samples were homogenized and total RNA was extracted using the RNeasy Plus Universal Mini Kit. RNA concentration and purity were assessed with a NanoDrop 2000 spectrophotometer (ThermoFisher Scientific). RNA integrity numbers (RIN) ranged from 8.3-9.3, as determined by Agilent TapeStation (Agilent Technologies). Library preparation was performed using the QIAseq Stranded mRNA Kit with 500ng total RNA. Library quality after purification was assessed using Agilent TapeStation D1000 (Agilent Technologies). High-quality libraries were pooled at equimolar concentrations and quantified using qPCR to ensure uniform sequencing representation. Sequencing was performed on a NextSeq 2000 (Illumina, San Diego, CA, USA) with a 1×75-cycle single-read and 2×10-cycle index run. Raw data were demultiplexed, generating individual FASTQ files using bcl2fastq v2.20.0.422 (Illumina). Sequencing data were analyzed using the Qiagen CLC Genomics Server 23.0.5. Ingenuity Pathway Analysis (IPA; Qiagen Bioinformatics, Redwood City, CA, USA) was used to identify enriched canonical pathways, upstream regulators, and interaction networks; upstream regulator symbols follow human-gene nomenclature as implemented in the IPA software (**S1.3**).

### Statistical analysis

If not otherwise noted, data were analyzed using GraphPad Prism 8.0 (GraphPad Software, San Diego, USA). For comparison of 2 experimental groups, 2-tailed Student’s t test with Welch’s correction was performed. More than 2 groups were compared using an ANOVA test with Bonferroni’s correction. The number of mice per experimental group was determined using G*Power software (Heinrich Heine University, Düsseldorf, Germany). Survival curves were estimated by the Kaplan-Meier method and compared using the log-rank (Mantel-Cox) test and the Gehan-Breslow-Wilcoxon test. Hazard ratios and corresponding 95% confidence intervals were calculated using the Mantel–Haenszel method. Unless stated otherwise, data are shown as mean ±SEM. A p-value < 0.05 was considered statistically significant.

## Results and discussion

### *In silico*-guided calculations defined conditions for a non-thermal, uniform acoustic exposure *in vitro*

At 1MHz, 1W/cm^2^, 100% duty cycle, *in silico* simulations (run until the emitted waves undergo several boundary reflections and establish a stable interference pattern) identified a 94mm transducer-to-sample distance as optimal for delivering a spatially homogeneous pressure field at the cell-culture plane while avoiding the localized maximum pressure that could cause hyperthermia (**Fig. 3A and S3**). Subsequent hydrophone measurements confirmed that, despite applying a non-focused ultrasound source, cells at this distance are exposed to a stable, spatially uniform pressure field (**Fig. 3B**; maximum pressure: 18.4kPa). Temperature measurements monitored inside a microplate during a 20min LIU treatment confirmed *in silico* predictions, showing that the temperature remained within the physiological range (36–39°C) under the chosen conditions (**Fig. 3C**).

**Fig. 3.**
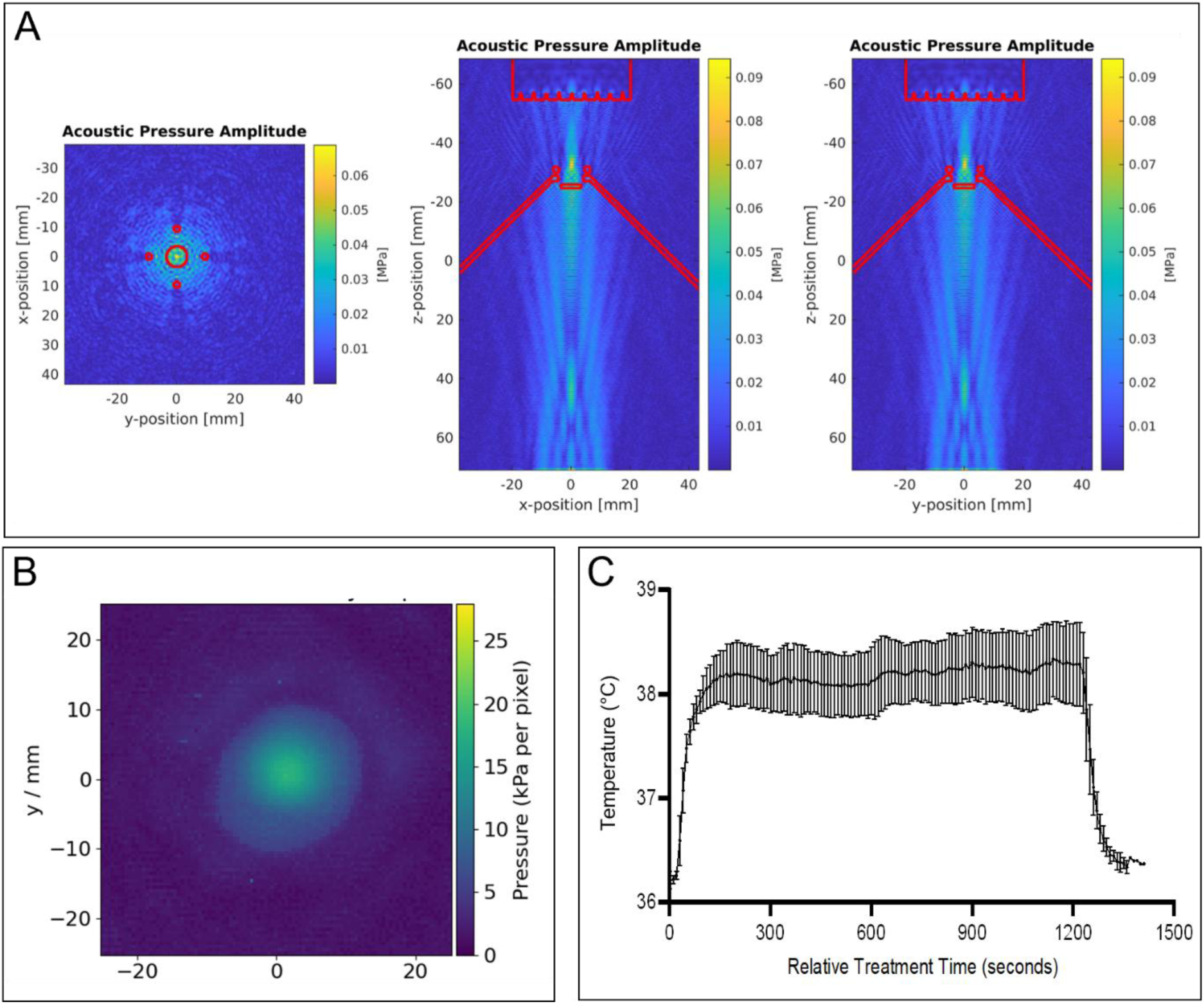
Acoustic field characterization and temperature stability during in vitro low-intensity ultrasound (LIU) exposure. (A) Simulated acoustic-pressure amplitude distribution at the optimized transducer-to-sample distance of 94mm. From left to right: transverse (x–y) section at the cell-culture plane, longitudinal x–z section through the center of the treatment geometry, and longitudinal y–z section through its center. The simulations demonstrate a spatially homogeneous pressure field at the cell-culture plane without localized pressure maxima. Red outlines indicate components of the modeled treatment geometry. (B) Hydrophone measurement of the pressure distribution at the cell-culture plane, confirming a stable and spatially uniform acoustic field, with a maximum measured pressure of 18.4kPa. (C) Continuous temperature monitoring inside the microplate during the 20min. LIU exposure. Temperature remained within the physiological range of 36–39°C, confirming non-hyperthermic exposure conditions. Simulations and experimental measurements were independently repeated at least three times. Data in (C) represent mean ± SEM.

### Exposure to LIU induces intracellular stress responses and apoptosis in 4T07 spheroids

*In silico*-defined LIU conditions, applied to 4T07 spheroids for 20min, resulted in structural damage, which was microscopically visible 24h post-treatment as less defined outer boundaries compared to untreated controls (**Fig. 4A**). Concomitantly, CellTiter-Glo assays demonstrated a statistically significant reduction in ATP levels, which decreased by more than 40% in treated spheroids compared to controls (**Fig. 4B**). As shown by flow cytometry, these functional changes were accompanied by significantly increased frequencies of Hsp70⁺, Hsp90⁺ and Annexin V⁺/Zombie UV⁺ cells in treated versus untreated spheroids (**Fig. 4C, D** and **S4**), collectively indicating LIU-induced intracellular stress, ongoing apoptosis, and metabolic impairment. While calreticulin and HMGB1, two canonical markers of immunogenic cell death, ICD [25], did not reach statistical significance under the examined conditions, it cannot be excluded that this may rather reflect the early 24h post-exposure timepoint, the spheroid geometry, or assay sensitivity. As shown below, our *in vivo* results point toward an involvement of ICD pathways. Future studies using surface calreticulin staining and HMGB1 quantification across multiple timepoints and in monolayer cultures would help clarify this. These findings are consistent with a prior study reporting analogous LIU-induced morphological and viability impairment in PANC-1 pancreatic cancer spheroids [26].

**Fig. 4.**
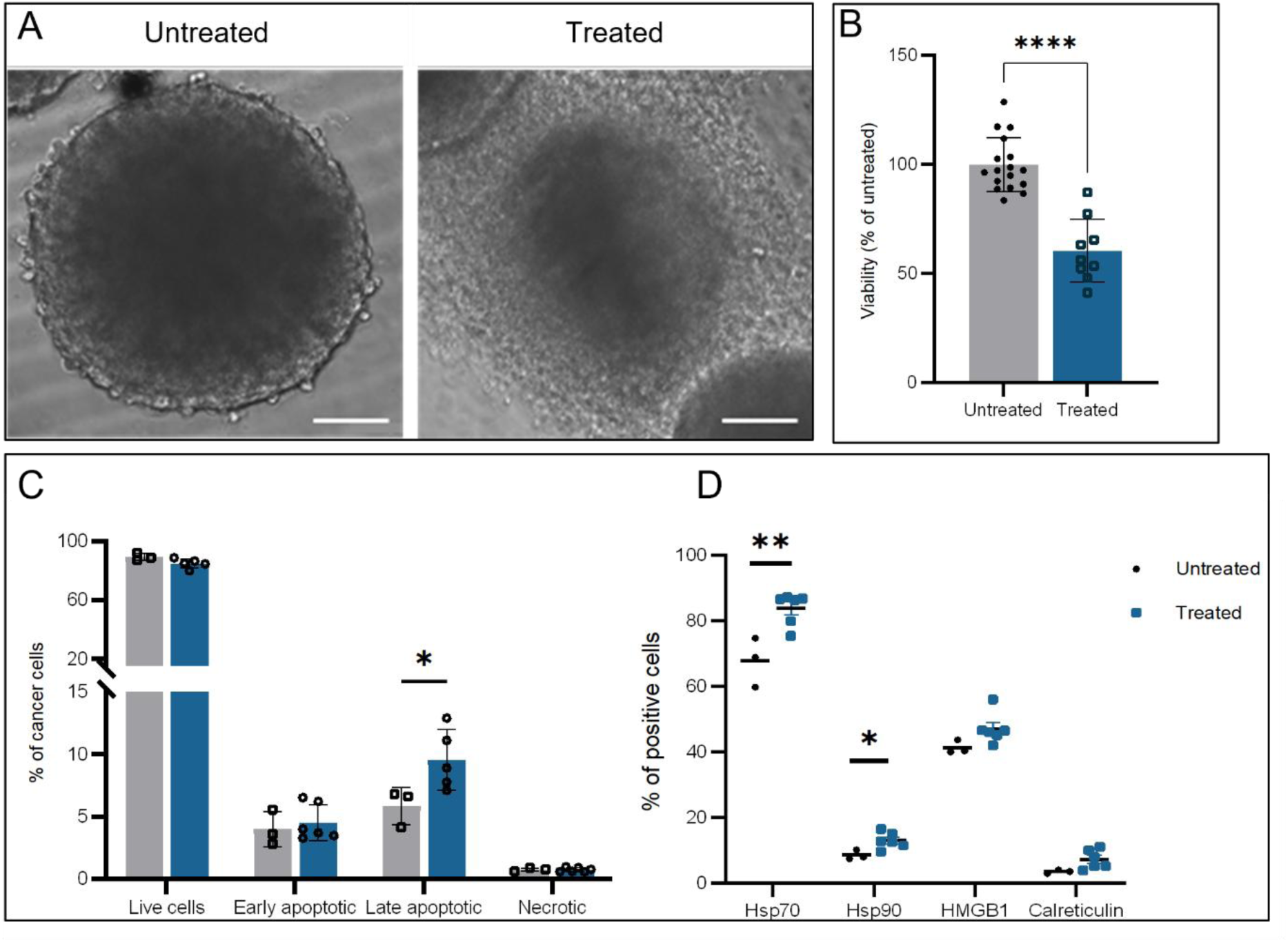
4T07 spheroid exposure to low-intensity ultrasound (LIU). (A) Representative brightfield images of untreated and LIU-treated (1MHz, 1W/cm^2^, 100% duty cycle, 20min.; “Treated”) spheroids 24h post-treatment. LIU-treated spheroids displayed disrupted morphology with less defined outer boundaries compared to controls. Scale bar: 200μm. (B) Cell viability assessed by the CellTiter-Glo 3D assay 24h after LIU exposure. Bars represent mean ± SEM; each data point represents one independent biological replicate in which 8 spheroids were pooled per condition. Data normalized to untreated controls. ****p < 0.0001 by unpaired t-test. (C) Flow cytometry analysis of Annexin V/Zombie UV staining at 24h after treatment revealed a significant increase in late apoptotic cells (Annexin V⁺/Zombie UV⁺) in LIU-treated spheroids compared to controls. Bars represent mean±SEM. *p<0.05 by unpaired t-test. (D) Flow cytometric quantification of four markers associated with immunogenic cell stress and death in spheroids 24 h after LIU exposure. A significant increase in Hsp70- and Hsp90-positive cells was detected in LIU-treated spheroids. Data represents the percentage of total cells positive for each marker, independent of Annexin V/Zombie UV status. *p<0.05, **p<0.01 by Welch’s t-test (Hsp90) or Mann-Whitney test (Hsp70). All experiments were independently repeated at least three times.

### Simulation-guided LIU setup enables safe and targeted *in vivo* ultrasound delivery

*In silico* simulations were performed to define an in vivo setup that concentrates acoustic exposure at the tumor site. Based on anatomically realistic computational mouse models and incorporating the complete treatment geometry, simulations predicted a relatively well-confined focal region within the tumor when the transducer is positioned 20mm above its surface, limiting energy deposition in adjacent, more acoustically absorptive internal organs (**Fig. 5A**). Thermal modeling based on tissue-specific acoustic and thermal properties indicated that this would cause neither hyperthermia within the tumor nor in the surrounding tissues, including those most susceptible to acoustic absorption (**Fig. 5A**). *In vivo* thermometry confirmed that intratumoral temperatures remained below 39°C throughout treatment (**Fig. 5B**), core body temperatures were stable, and no macroscopic skin lesions were observed in any animal, collectively confirming the safety and tolerability of the protocol.

**Fig. 5.**
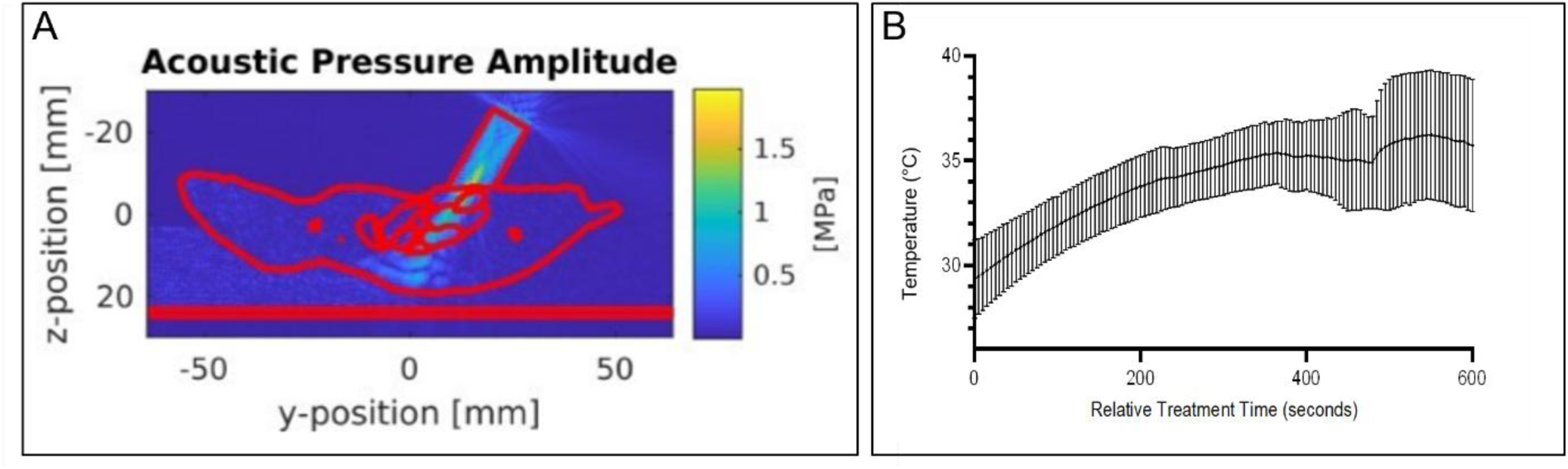
Simulation-guided low-intensity ultrasound (LIU) setup and intratumoral temperature monitoring. (A) *In silico* simulation of the acoustic pressure field generated by the transducer at the tumor site. As expected for an interference pattern, a peak amplitude occurred close to the skin boundary; because the tumor was located superficially, this peak coincided with the tumor region rather than extending into deeper, more acoustically absorptive organs. (B) Intratumoral temperature monitoring during LIU exposure. Real-time recordings show a gradual increase in temperature that remained below 39°C throughout the 10min. treatment, confirming non-thermal conditions under the applied LIU protocol.

### Cyclic LIU treatment delays tumor growth and improves survival *in vivo*

All mice injected with 4T07 cells developed a palpable tumor, reaching a minimum volume of 100mm^3^ within 7 days. Mice of the treatment group were then treated with LIU (1MHz, 1W/cm^2^, 100% duty cycle, 10min per session) every third day for up to 6 treatment cycles, with a 20mm gel pad placed between the transducer and the tumor site to maintain the simulated optimal transducer-to-tumor distance (**Fig. 2B**). Following the first and second LIU cycle, tumor volumes in the treated group were significantly smaller than in untreated animals (**Fig. 6A, C**). Due to an increasing number of untreated animals reaching the ethical withdrawal criterion (tumor volume ≥1000mm^3^), cross-sectional tumor-volume comparisons beyond the second cycle were not feasible. However, time-to-endpoint data from all animals were included in the Kaplan-Meier survival analysis. Over the full 6-cycle observation period, this revealed a clear divergence in survival probability between groups (**Fig. 6B**), confirmed by both the log-rank (p = 0.0008) and Gehan-Breslow-Wilcoxon (p = 0.0007) tests. Median survival increased from 3 cycles in the untreated group to 4 cycles in the LIU-treated group. The Mantel-Haenszel hazard ratio was 0.09 (95% CI: 0.02–0.37); the wide confidence interval reflects the reduced effective sample size at later timepoints, driven by rapid tumor progression in untreated animals. These results support a genuine survival benefit of LIU treatment and provide a strong basis for follow-up studies.

**Fig. 6.**
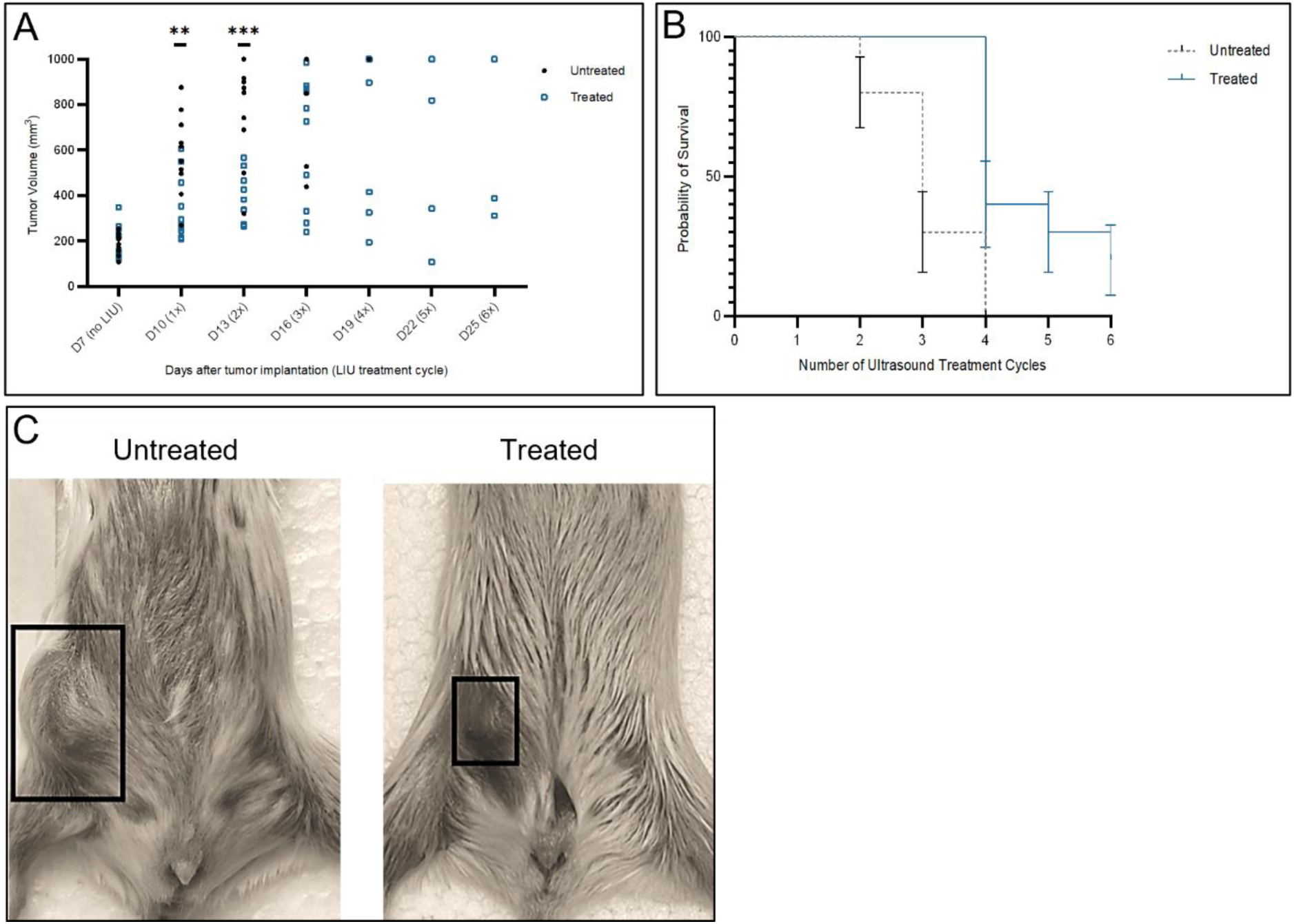
Low-intensity ultrasound (LIU)-mediated effects on tumor volume and survival *in vivo.* BALB/c mice bearing orthotopic 4T07 mammary tumors were subjected to repeated LIU treatments (per cycle: 1MHz, 1W/cm^2^, 100% duty cycle, 10min; up to six cycles, 1x-6x) under non-thermal conditions. (A) Tumor growth curves showing a significant reduction in tumor volume in LIU-treated mice compared with untreated controls after the first and second treatment cycles (**p≤0.01; ***p≤0.001). Error bars represent mean±SEM. (B) Kaplan–Meier survival analysis demonstrating improved survival in LIU-treated mice across six treatment cycles (Log-rank [Mantel–Cox] test, ***p<0.001). (C) Representative images of untreated and LIU-treated mice after six ultrasound cycles, showing visibly reduced tumor size, outlined by black boxes.

Prior continuous LIU studies have been largely confined to *in vitro* systems or drug-delivery applications with combination agents [27]. Non-ablative pulsed focused ultrasound has been shown to delay tumor growth in syngeneic murine models [28,29], and spleen-targeted pulsed LIU has demonstrated immune-mediated growth inhibition in 4T1 mammary tumors [30]. To our knowledge, the present study is the first to demonstrate standalone antitumor efficacy of continuously applied LIU delivered to the tumor in an orthotopic syngeneic breast cancer model. The one-cycle improvement in median survival is consistent with non-ablative ultrasound acting as an immune-priming intervention, whose efficacy may benefit from combinations with immune checkpoint blockade [6,30].

### A single LIU cycle induces acute tumor necrosis and innate immune cell infiltration

To characterize the immediate tissue-level consequences of LIU, tumors were harvested 24h after the first treatment cycle. Quantitative analysis of H&E-stained sections revealed that the percentage of necrotic areas relative to the total tumor area was significantly greater in LIU-treated tumors compared to untreated controls (**Fig. 7**), demonstrating that a single LIU session is sufficient to induce measurable tumor destruction *in vivo*. Immunohistochemical analysis revealed concurrent changes in the immune composition of TME. Iba1^+^ macrophages were significantly elevated in treated tumors, with focal enrichment in necrotic and peritumoral regions, while CD45R^+^ B cells were also significantly increased and distributed diffusely throughout the TME (**Fig. 8**). The disproportionately larger Iba1^+^ tissue area relative to CD45R^+^ area likely reflects the branching morphology of macrophages, which occupy greater area per cell than the compact B cells presumed to numerically dominate the CD45R^+^ compartment. In contrast, CD3^+^, CD4^+^, and CD8^+^ T-cell infiltration did not differ significantly between groups at this early post-treatment time point (**Fig. 8**).

**Fig. 7.**
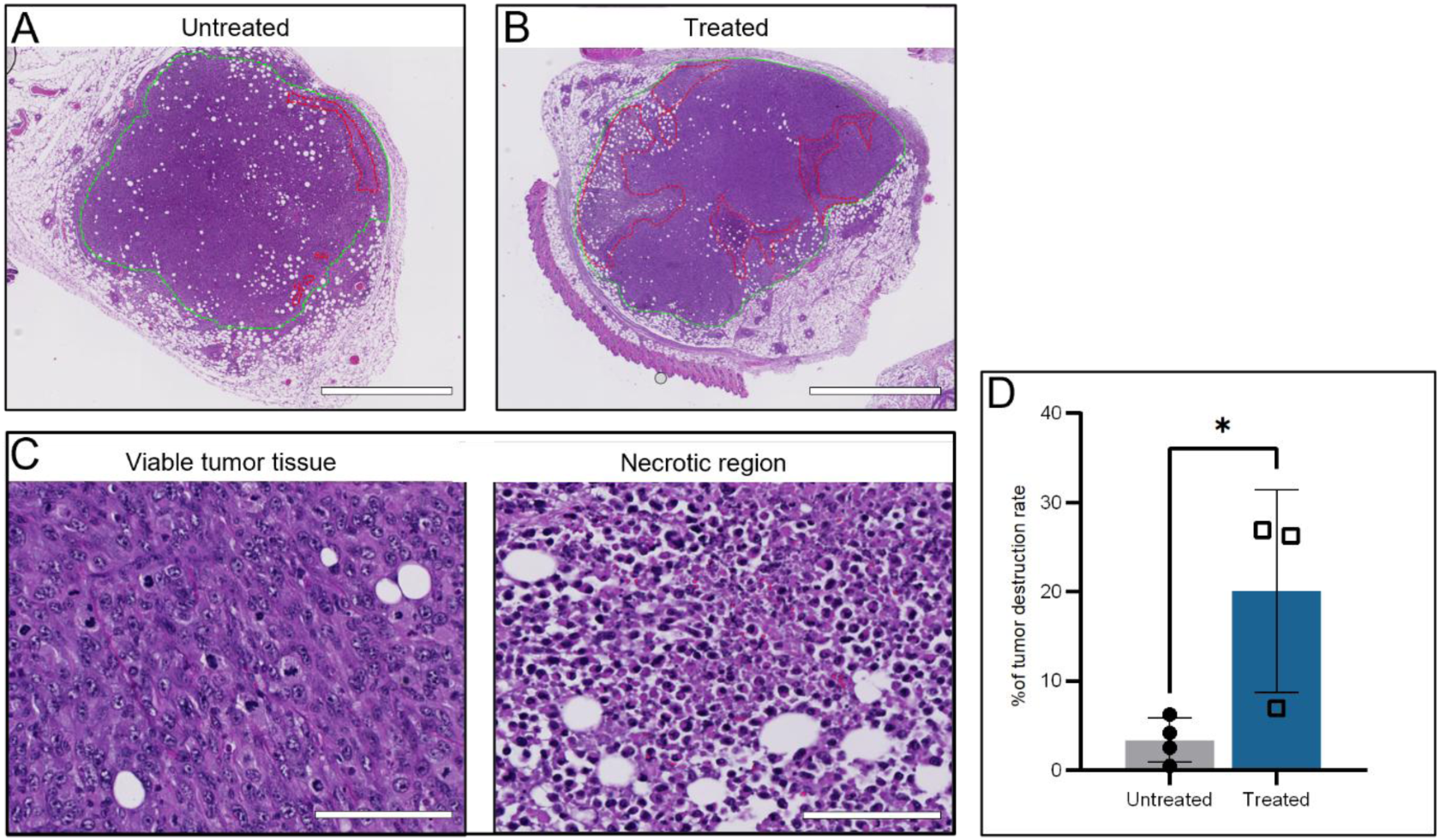
Histopathological changes in 4T07 tumors. Tumor samples were collected 24h after the first cycle of LIU treatment (1MHz, 1W/cm^2^, 100% duty cycle, 10min.; “Treated”) and compared to untreated controls (“Untreated”) after Hematoxylin and eosin (H&E) staining. (A–B) Representative low-magnification images of untreated (A) and LIU-treated (B) tumors. The total tumor area is outlined in green, and necrotic regions are outlined in red. Scale bars: 2mm. (C) High-magnification views of viable tumor tissue (left) and necrotic region (right) in LIU-treated tumors. Scale bars: 100µm. (D) Quantification of the tumor destruction rate, expressed as the percentage of necrotic area relative to total tumor area, revealed significantly increased necrosis in LIU-treated tumors compared with untreated controls (p<0.05, unpaired t-test). Bars represent mean ± SD; individual data points denote biological replicates.

**Fig. 8.**
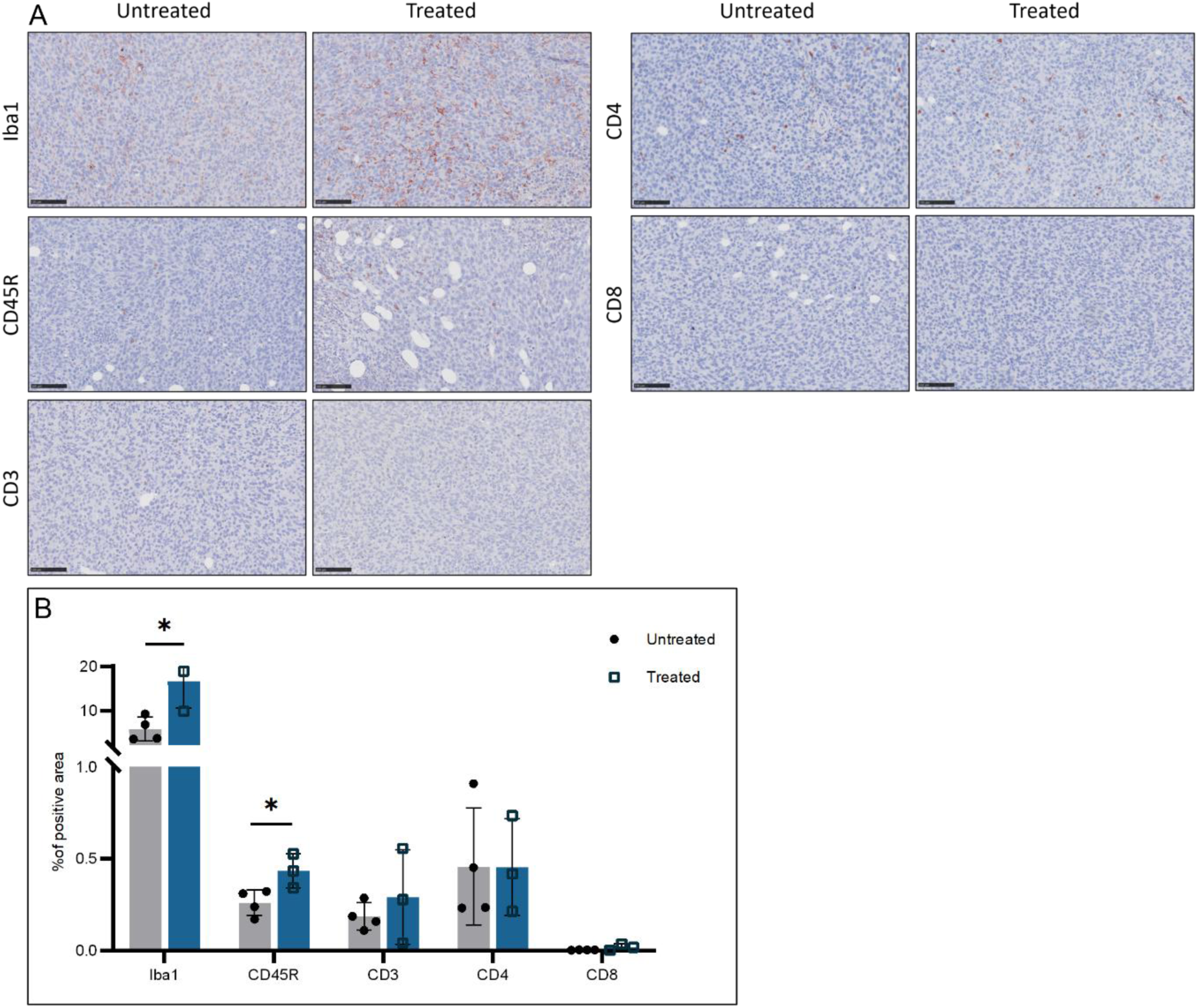
Immunohistochemistry of immune cell infiltration in 4T07 tumors. (A) Representative images of 4T07 tumors 24h after the first LIU cycle (1MHz, 1W/cm^2^, 100% duty cycle, 10min.) compared to untreated controls. Sections were stained for Iba1 (macrophages), CD45R (B cells), CD3 (pan-T cells), CD4 (helper T cells), and CD8 (cytotoxic T cells). AEC chromogen (red) was used for detection, with hematoxylin counterstaining (blue). Scale bars: 100µm. (B) Quantification of Iba1-, CD45R-, CD3-, CD4-, and CD8-positive areas expressed as the percentage of total tumor area (% positive area) determined by digital pathology analysis. Bars represent mean±SD; dots indicate biological replicates. p<0.05 by unpaired t-test.

The pattern of early myeloid recruitment without concurrent T-cell infiltration is consistent with the kinetics of innate immune activation following early tissue damage. Similar innate-predominant infiltration has been reported following pulsed and focused ultrasound in syngeneic tumor models [28,31], suggesting this may be a shared feature of physical ultrasound modalities independent of sonosensitizing agents. These early findings suggest that LIU initiates a pro-inflammatory cascade in the TME, predominantly through innate immune activation, which may create conditions favorable for subsequent adaptive immune engagement.

### Repeated LIU cycles drive broad transcriptional reprogramming of the TME

This study reports, for the first time, whole-transcriptome profiling of tumor tissue following continuous LIU treatment in a syngeneic *in vivo* model. To capture transcriptional changes following repeated exposure, bulk RNA sequencing was performed on tumors collected 24h after the third treatment cycle. Principal component analysis (PCA) of the 500 most variable genes revealed a clear separation between LIU-treated and untreated samples (**Fig. 9A**), confirmed by hierarchical clustering (**Fig. 9B**). Functional analysis identified three major clusters related to (1) immune cell proliferation, activation, and migration, (2) non-immune cell growth and progression, and (3) inflammatory responses and cytokine production. A comprehensive gene list is provided in **Table S1**.

**Fig. 9.**
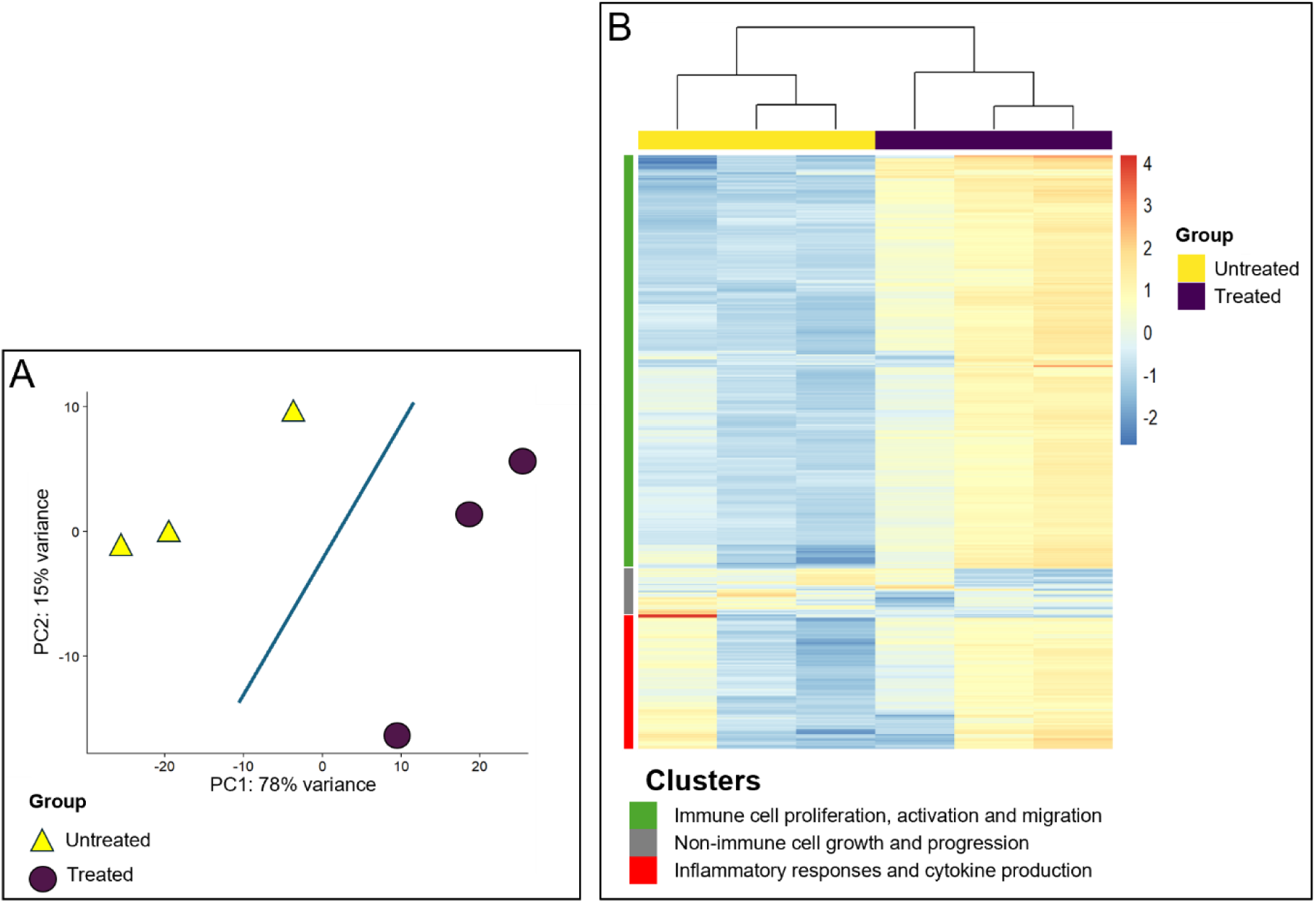
Transcriptional changes in 4T07 tumors. Exploratory analysis of distribution and homogeneity in 4T07 tumors 24h after the third LIU cycle (1MHz, 1W/cm^2^, 100% duty cycle, 10min.; “Treated”) compared to untreated controls (“Untreated”), considering the 500 most variable genes among all samples. (A) Principal component (PC) analysis plot of all samples, illustrating a clear separation between experimental groups. (B) Heatmap of the 500 genes varying the most between all samples. Colors yellow to violet indicate a gradient of high to low expression of each gene related to its average expression. LIU-treated and untreated samples form distinct clusters, confirming LIU-driven transcriptional reprogramming.

Out of 23,664 expressed genes, 2,573 differentially expressed genes (DEGs) met significance thresholds (p≤0.05, FDR≤0.05), comprising 2,245 upregulated and 328 downregulated genes (**Fig. 10**). Gene ontology (GO) analysis of upregulated DEGs revealed enrichment in immune activation (regulation of cytokine production, p=5.03×10^-18^; T-cell activation, p=1.78×10^-15^; B-cell activation, p=2.09×10^-9^), inflammatory signaling (inflammatory response, p=3.26×10^-13^), and tissue remodeling (extracellular matrix, ECM, organization, p=2.18×10^-7^), among others (**Fig. 10**; **Supplementary File S1**). Downregulated DEGs were enriched for tumor-intrinsic proliferative programs, including regulation of cell population proliferation (p=6.92×10^-5^), response to steroid hormone (p=9.89×10^-5^), and mammary gland development (p=4.11×10^-4^) (**Fig. 10**). Network analysis confirmed that only upregulated DEGs formed significant functional networks, centered on immune regulation, cytokine production, ECM organization, and cell death pathways (**Fig. 11**; **Table S1**).

**Fig. 10.**
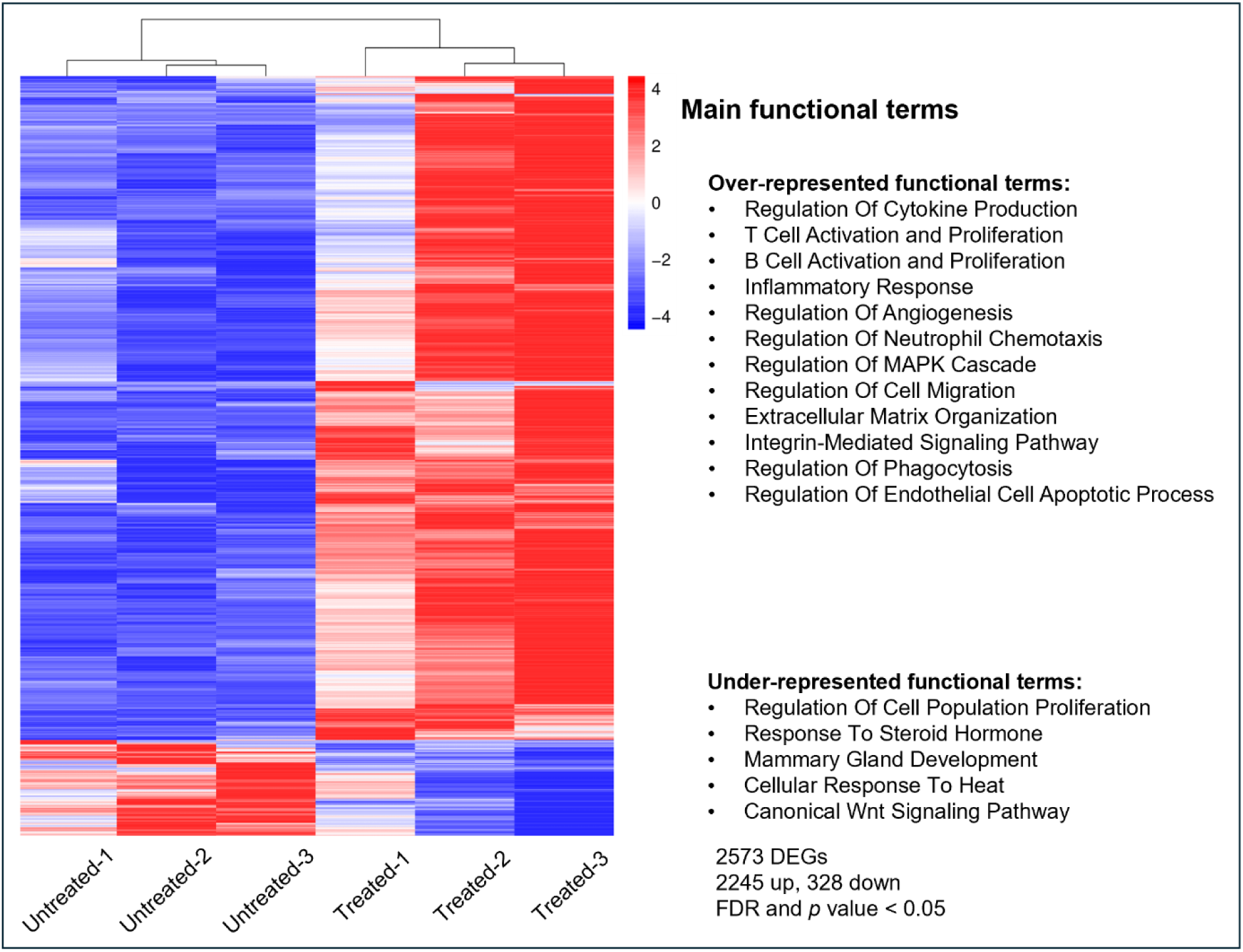
Heatmap and overrepresented gene ontologies. The analysis revealed a significant transcriptional difference between tumor samples collected after the third cycle of low-intensity ultrasound (1MHz, 1W/cm^2^, 100% duty cycle, 10 min.; “Treated”) and from the untreated group (“Untreated”; p<0.05, FDR<0.05), with an upregulation of 2245 differentially expressed genes (DEGs), and downregulation of 328 DEGs. The expression gradient of each gene, relative to its abundance (log2 ratio), is depicted by colors ranging from red to blue. The main over- and underrepresented biological processes and gene ontologies indicated were identified using Enrichr.

**Fig. 11.**
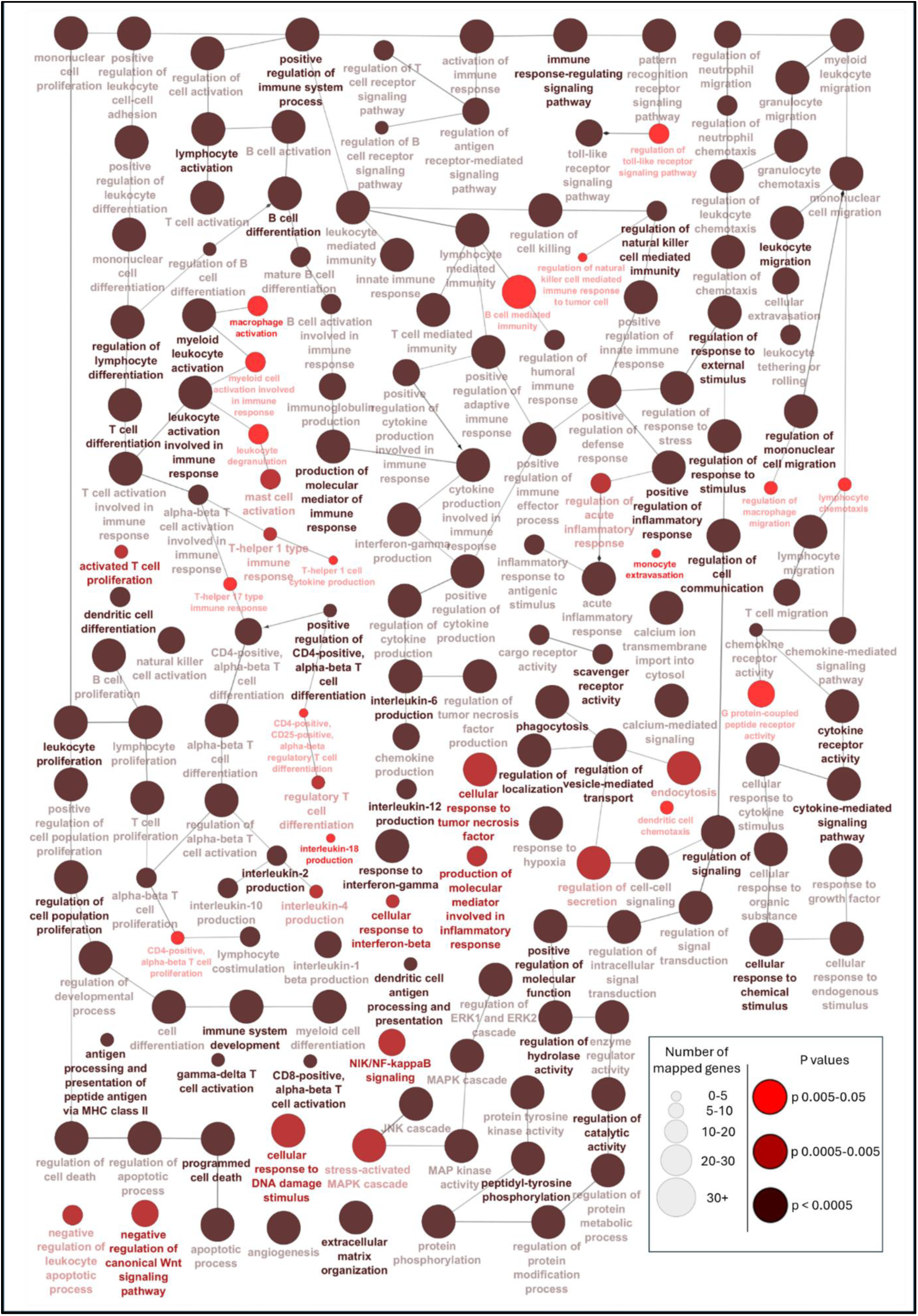
Functional networks for differentially expressed genes (DEGs) in tumors treated with low-intensity ultrasound (US) vs. untreated tumors. Over-represented networks were determined using ClueGO from Cytoscape software. Networks were manually rearranged and narrowed down by p<0.05, excluding redundant and non-informative terms. The number of mapped genes for each biological term is represented by node size, statistical significance is denoted by node colour (see legend). Networks most strongly represented among both up- and downregulated DEGs after US were associated with: immune system activation, T-cell and B-cell differentiation and activation, regulation of cytokine production, extracellular matrix organization, regulation of cell population proliferation, apoptosis, regulation of cell signaling, and angiogenesis.

IPA identified broad activation of immune and inflammatory canonical pathways, most prominently Th1 (p=2.00×10^-24^) and Th2 (p=3.16×10^-23^) signaling, macrophage classical (p=6.31×10^-11^) and alternative (p=2.51×10^-17^) activation, T-cell receptor signaling (p=8.32×10^-6^), and IFN-γ signaling (p=6.17×10^-8^). The predominance of classical over alternative macrophage activation pathway enrichment is consistent with a shift toward M1 polarization, one of the most consistently reported effects of mechanical ultrasound in preclinical tumor models [28,31]. Immunosuppressive pathways were concurrently deactivated, including IL-10 signaling (p=6.46×10^-10^) and MSP-RON signaling in macrophages (p=2.75×10^-6^). Upstream regulator analysis identified TNF, IFN-γ, IL-2, IL-4, IL-1β, and IL-6 as positively activated, consistent with broad pro-inflammatory cytokine signaling. The complete list of canonical pathways, upstream regulators, and associated DEGs is provided in **Table S1** and **Supplementary File S1**. As bulk RNA-seq cannot distinguish transcriptional reprogramming from immune cell compositional shifts, the strong DEG asymmetry (87% upregulated, predominantly immune genes) is interpreted as hypothesis-generating.

### LIU-associated changes in transcriptional regulation of tumor growth and progression

Consistent with reduced tumor burden *in vivo*, LIU downregulated a coherent set of genes driving tumor growth, angiogenesis, and metastatic dissemination, including matrix metalloproteinases (*Mmp1a*, *Mmp10*), tumor-stroma signaling factors (*Tgfa*, *Tgfb3*, *Pdgfa*), and additional pro-tumorigenic mediators (*Nupr1*, *Hspa1a*, *Sqle*, *Il11*, *Wap*, *Saa1*). Conversely, LIU upregulated genes associated with tumor suppression and pyroptosis, most notably *Pdcd4*, *Casp4*, and *Gsdmd* [32–34], indicating a concerted transcriptional shift away from pro-tumorigenic programs and toward inflammatory cell-death pathways (**Supplementary File S1**). Mechanistically, suppression of *Mmp1a* and *Mmp10* implies reduced ECM degradation and local invasion capacity, while lower *Tgfa* expression suggests attenuation of tumor-stroma signaling and weakening of an established immune-evasion axis [35,36].

### LIU-associated transcriptional changes in immunogenic and inflammatory pathways

Repeated LIU treatment was associated with upregulation of genes linked to ICD (*Ifngr1*, *Nlrp1b*, *Il1b*, *Tnf*, *Pdcd4*, *Cxcl11*) and broad engagement of innate pattern-recognition receptor pathways, including multiple Toll-like receptor family members (*Tlr1*, *Tlr2*, *Tlr7*–*Tlr9*, *Tlr11*–*Tlr13*), C-type lectin receptors (*Clec4a1*, *Clec9a*), and inflammasome components (*Nlrp3*, *Casp4*, *Mefv*, *Gsdmd*), indicating coordinated activation of danger-sensing and sterile inflammatory programs (**Supplementary File S1**). NLRP3 inflammasome activation can promote dendritic cell maturation and CD8^+^ T-cell priming [37,38]; the concurrent upregulation of *Nlrp3*, *Casp4* and *Gsdmd* transcripts observed is consistent with engagement of this pathway. Upregulation of *Cd36* further reinforces sterile inflammatory signaling through ROS generation and pro-inflammatory cytokine production [39].

### LIU-associated transcriptional changes related to innate and adaptive immune responses

Innate immune sensing was enhanced by upregulation of pattern-recognition receptors (*Tlr2*, *Nod1*, *Nod2*, *Clec4a1*, among others) and chemoattractants (*Cxcl5*, *Cxcl9*, *Cxcl11*, *Ccl21a*), consistent with enhanced recruitment of myeloid cells and dendritic cells to the TME (**Supplementary File S1**). Upregulation of T-cell-associated genes (*Cd3d*, *Cd3e*, *Cd3g*, *Cd4*, *Cd8a*, *Cd8b1*, *Cd69*, *Il2ra*) and costimulatory/checkpoint molecules (*Tnfrsf4*, *Icos*, *Ctla4*) was observed. However, protein-level validation of T-cell infiltration at the cycle-3 timepoint was not performed, so this transcriptomic signal, while consistent with a progression from early innate responses toward later lymphocyte engagement reported for other immunomodulatory physical stimuli [28], may partly reflect changes in immune-cell composition rather than *de novo* activation. LIU additionally upregulated *Cd19*, *Ms4a1*, *Cd27*, *Blk*, and *Cxcl13*, pointing to engagement of the B-cell compartment, though whether this extends to durable immune memory remains to be established. To our knowledge, transcriptional engagement of the B-cell compartment has not previously been reported for standalone continuous mechanical ultrasound. This transcriptional signal is consistent with the protein-level increase in CD45R^+^ (B220^+^) cells observed by immunohistochemistry 24h after a single LIU cycle (**Fig. 8**), suggesting that B-cell engagement begins early and is sustained, or amplified, with repeated treatment. The concurrent predicted inhibition of MSP-RON signaling and IL-10 pathway deactivation further suggests that LIU actively dismantles specific immunosuppressive programs rather than broadly amplifying immune activity.

Collectively, the transcriptomic landscape after three LIU cycles depicts a TME undergoing coordinated reprogramming, away from immunosuppressive and pro-tumorigenic programs and toward innate activation, danger signaling, and early adaptive immune engagement, providing a molecular basis for the antitumor efficacy observed *in vivo*.

### Repeated LIU cycles reverse splenomegaly and modulate systemic cytokine profiles

Untreated tumor-bearing mice developed progressive splenomegaly relative to healthy controls, consistent with previous reports linking splenomegaly in 4T07 and related mammary tumor models to tumor-driven myelopoiesis and systemic expansion of myeloid-derived suppressor cells (MDSCs) [40–42]. To our knowledge, we show here for the first time that continuous LIU can significantly reduce tumor-associated splenomegaly (**Fig. 12A**), with LIU-treated mice remaining closer to healthy spleen volumes from cycle 3 onward. This raises the possibility that splenic regression could serve as a surrogate marker of therapeutic efficacy, an association reported in other cancer models and in patients receiving immunotherapy [43–45], though formal correlation with tumor outcome in this model remains to be tested.

**Fig. 12.**
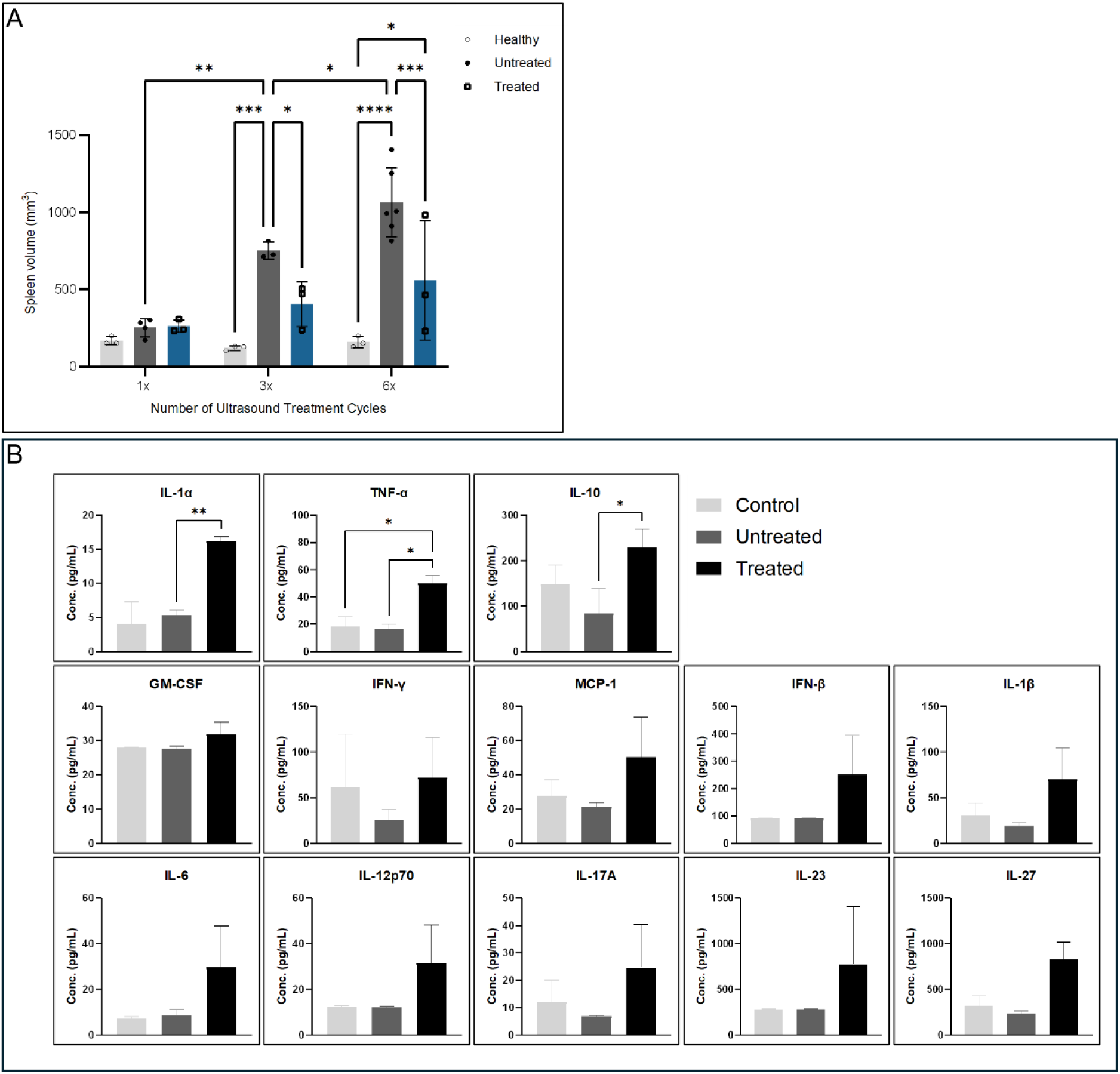
Spleen volumes and systemic cytokine profiles. Effects were measured in healthy, 4T07 tumor-bearing untreated, and 4T07 tumor-bearing mice treated with low-intensity ultrasound (LIU; 1MHz, 1W/cm^2^, 100% duty cycle, 10min.) (A) Spleen volumes (mm^3^) were measured after one (1x), three (3x) and six (6x) treatment cycles in 4T07 tumor-bearing mice (“Treated”), compared to the respective time points in untreated 4T07 tumor-bearing mice (“Untreated”) and tumor-free controls (“Healthy”). Error bars represent mean±SEM. *p<0.05; **p<0.01; ***p<0.001; ****p<0.0001. (B) Plasma cytokine concentrations (pg/mL) measured 24h after three LIU treatment cycles. Data are presented as mean±SEM (\**p*<0.05, \*\**p*<0.01, \*\*\**p*<0.001).

To determine whether the attenuation of splenomegaly was accompanied by systemic immune changes, plasma cytokine concentrations were measured 24h after the third LIU cycle. Using a multiplex cytokine immunoassay (**Fig. 12B**), we found that IL-1α, TNF-α, and IL-10 were significantly elevated in LIU-treated compared to untreated tumor-bearing mice, while levels of IFN-γ, IFN-β, IL-1β, IL-6, IL-12p70, IL-17A, IL-23, IL-27, MCP-1 and GM-CSF did not reach statistical significance. The significant elevation of IL-1α and TNF-α is consistent with activation of innate inflammatory pathways that can enhance antigen presentation and leukocyte recruitment [46,47]. However, IL-1α has also been shown to drive context-dependent immunosuppressive effects within the TME by reprogramming tumor-associated myeloid cells [48]. The concurrent significant elevation of IL-10 introduces further complexity: as a pleiotropic immunoregulatory cytokine, IL-10 can suppress Th1 polarization and cytotoxic T-cell activity, while limiting excessive inflammatory tissue damage [49]. The simultaneous elevation may therefore reflect a dynamic balance between inflammatory activation and compensatory immunoregulation.

## Summary and conclusions

This study set out to determine whether a controlled, non-thermal continuous LIU exposure can induce immunogenic stress responses, remodel tumor architecture, reprogram the TME toward an antitumor immune state, and elicit systemic immune responses in a syngeneic orthotopic model of breast cancer.

*In vitro*, a single 20min LIU exposure reduced metabolic activity in 4T07 tumor spheroids and induced intracellular stress responses and apoptosis. *In vivo*, cyclic LIU treatment significantly increased intratumoral necrosis, slowed tumor growth, and prolonged time to ethical endpoint compared with untreated controls, while whole-tumor RNA sequencing and immunohistochemistry showed that LIU reprograms the TME toward an antitumor state, with early myeloid and CD45R^+^ B-cell infiltration at cycle 1 followed by M1 macrophage, B-cell, and T-cell transcriptional signatures by cycle 3. These local effects were accompanied by systemic immune changes, including reversal of tumor-driven splenomegaly and significant elevation of plasma IL-1α, TNF-α, and IL-10.

To our knowledge, this is the first study to demonstrate that continuous, non-thermal LIU, without pharmacological or cavitation-enhancing agents, can drive both local tumor destruction and systemic antitumor immune engagement *in vivo*, including transcriptional and protein-level evidence of B-cell compartment engagement not previously reported for mechanical ultrasound alone.

These findings should be interpreted alongside limitations that define priorities for follow-up work. Animal numbers were set by power calculations for the primary endpoint (*i.e*., tumor volume/survival), leaving relatively small replicate numbers for the different analytical methods (*e.g.,* RNA-seq, cytokine assays, histopathology). Our secondary findings are accordingly hypothesis-generating, pending protein-level validation, larger sample sizes and additional time points.

Collectively, our findings establish a reproducible preclinical platform and a tractable mechanistic hypothesis centered on myeloid-driven innate immune remodeling and progressive engagement of the adaptive, including B-cell, compartment. Its non-invasive, drug-free nature may position LIU as a potential complement for immune checkpoint blockade and other immunotherapies.

## Acknowledgements

We sincerely thank Serkan Sariyildiz and Theresa Lehmann (Institute of Anatomy, University of Zurich) for their technical support and the Laboratory Animal Services Center (LASC) and Zurich Integrative Rodent Physiology (ZIRP), University of Zurich, for their support with animal husbandry and related procedures. We are grateful to Maries van den Broek and Paulo Pereira, Institute of Experimental Immunology, University of Zurich, Switzerland for helpful discussions and the provision of the 4T07 cell line.

## Funding

This work was supported by grants from Eureka Eurostars (E!114157) and the Karl & Rena Theiler-Haag Stiftung.

## Conflict of interests

The authors declare that they have no known competing financial interests or personal relationships that could have appeared to influence the work reported in this paper.

## CRediT authorship contribution statement

**Reyhaneh Hooshmandabbasi:** Conceptualization, Project administration, Supervision, Funding acquisition, Investigation, Methodology, Formal analysis, Data curation and integration, Validation, Visualization, Writing – original draft, Writing – review & editing. **Ali Kazemian:** Investigation, Methodology, Formal analysis (bioinformatics), Visualization, Writing – original draft, Writing – original draft, Writing – review & editing. **Rahul Singha:** Investigation, Methodology, Formal analysis, Validation, Visualization, Writing – original draft, Writing – review & editing. **Manuel Vielma Blanco:** Methodology, Software, Formal analysis, Visualization, Validation, Writing – original draft, Writing – review & editing. **Niloufar Nikkhah Bahrami:** Methodology, Formal analysis, Visualization, Writing – original draft, Writing – review & editing. **Thomas Hauser:** Methodology, Formal analysis, Visualization, Validation, Writing – review & editing. **Mathias Weyland:** Methodology, Formal analysis, Visualization, Validation, Writing – original draft, Writing – review & editing. **Franco Guscetti:** Methodology, Writing – review & editing. **Daniel Fehr:** Methodology, Investigation, Writing – original draft. **David Wahl:** Conceptualization, Supervision, Funding acquisition, Writing – review & editing. **Mathias Bonmarin:** Methodology, Writing – review & editing. **Stephan Scheidegger:** Methodology, Writing – review & editing. **Caroline Maake:** Conceptualization, Project administration, Supervision, Funding acquisition, Resources, Writing – review & editing.

## Data availability

The data that support the findings of this study are available on request from the corresponding author.

## Declaration of generative AI and AI-assisted technologies in the manuscript preparation process

During the preparation of the Graphical Abstract, the authors used BioRender (BioRender.com) and ChatGPT (OpenAI) to generate the graphical abstract, including some icon elements. After using these tools, the authors reviewed and edited the content as needed and take full responsibility for the content of the published article.

## Notes

### Competing Interest Statement

The authors have declared no competing interest.

## References

[1] Bray F, Laversanne M, Sung H, Ferlay J, Siegel RL, Soerjomataram I, et al. Global cancer statistics 2022: GLOBOCAN estimates of incidence and mortality worldwide for 36 cancers in 185 countries. CA A Cancer J Clinicians 2024;74(3):229–63. 10.3322/caac.21834.

[2] Hanahan D. Hallmarks of Cancer: New Dimensions. Cancer Discovery 2022;12(1):31–46. 10.1158/2159-8290.CD-21-1059.

[3] Waks AG, Winer EP. Breast Cancer Treatment. JAMA 2019;321(3):288. 10.1001/jama.2018.19323.

[4] Ribas A, Wolchok JD. Cancer immunotherapy using checkpoint blockade. Science 2018;359(6382):1350–5. 10.1126/science.aar4060.

[5] Galon J, Bruni D. Approaches to treat immune hot, altered and cold tumours with combination immunotherapies. Nat Rev Drug Discov 2019;18(3):197–218. 10.1038/s41573-018-0007-y.

[6] Sheybani ND, Price RJ. Perspectives on Recent Progress in Focused Ultrasound Immunotherapy. Theranostics 2019;9(25):7749–58. 10.7150/thno.37131.

[7] Bader KB, Padilla F, Haworth KJ, Ellens N, Dalecki D, Miller DL et al. Overview of Therapeutic Ultrasound Applications and Safety Considerations: 2024 Update. J of Ultrasound Medicine 2025;44(3):381–433. 10.1002/jum.16611.

[8] Izadifar Z, Izadifar Z, Chapman D, Babyn P. An Introduction to High Intensity Focused Ultrasound: Systematic Review on Principles, Devices, and Clinical Applications. JCM 2020;9(2):460. 10.3390/jcm9020460.

[9] Izadifar Z, Babyn P, Chapman D. Mechanical and Biological Effects of Ultrasound: A Review of Present Knowledge. Ultrasound in Medicine & Biology 2017;43(6):1085–104. 10.1016/j.ultrasmedbio.2017.01.023.

[10] Skalina KA, Singh S, Chavez CG, Macian F, Guha C. Low Intensity Focused Ultrasound (LOFU)-mediated Acoustic Immune Priming and Ablative Radiation Therapy for in situ Tumor Vaccines. Sci Rep 2019;9(1). 10.1038/s41598-019-51332-4.

[11] van den Bijgaart RJE, Eikelenboom DC, Hoogenboom M, Fütterer JJ, Brok MH den, Adema GJ. Thermal and mechanical high-intensity focused ultrasound: perspectives on tumor ablation, immune effects and combination strategies. Cancer Immunol Immunother 2017;66(2):247–58. 10.1007/s00262-016-1891-9.

[12] Tardoski S, Ngo J, Gineyts E, Roux J-P, Clézardin P, Melodelima D. Low-intensity continuous ultrasound triggers effective bisphosphonate anticancer activity in breast cancer. Sci Rep 2015;5(1). 10.1038/srep16354.

[13] Bazou D, Maimon N, Munn L, Gonzalez I. Effects of Low Intensity Continuous Ultrasound (LICU) on Mouse Pancreatic Tumor Explants. Applied Sciences 2017;7(12):1275. 10.3390/app7121275.

[14] Kim MG, Yoon C, Lim HG. Recent Advancements in High-Frequency Ultrasound Applications from Imaging to Microbeam Stimulation. Sensors 2024;24(19):6471. 10.3390/s24196471.

[15] Gong Z, Dai Z. Design and Challenges of Sonodynamic Therapy System for Cancer Theranostics: From Equipment to Sensitizers. Advanced Science 2021;8(10). 10.1002/advs.202002178.

[16] Araújo Martins Y, Zeferino Pavan T, Fonseca Vianna Lopez R. Sonodynamic therapy: Ultrasound parameters and in vitro experimental configurations. International Journal of Pharmaceutics 2021;610:121243. 10.1016/j.ijpharm.2021.121243.

[17] Frutos Díaz-Alejo J, Gonzalez Gomez I, Earl J. Ultrasounds in cancer therapy: A summary of their use and unexplored potential. Oncol Rev 2022;16(1). 10.4081/oncol.2022.531.

[18] Jenkins L, Jungwirth U, Avgustinova A, Iravani M, Mills A, Haider S et al. Cancer-Associated Fibroblasts Suppress CD8+ T-cell Infiltration and Confer Resistance to Immune-Checkpoint Blockade. Cancer Research 2022;82(16):2904–17. 10.1158/0008-5472.CAN-21-4141.

[19] Miller BE, Miller FR, Wilburn DJ, Heppner GH. Analysis of tumour cell composition in tumours composed of paired mixtures of mammary tumour cell lines. Br J Cancer 1987;56(5):561–9. 10.1038/bjc.1987.242.

[20] Treeby BE, Cox BT. k-Wave: MATLAB toolbox for the simulation and reconstruction of photoacoustic wave fields. J Biomed Opt 2010;15(2):21314. 10.1117/1.3360308.

[21] Treeby BE, Budisky J, Wise ES, Jaros J, Cox BT. Rapid calculation of acoustic fields from arbitrary continuous-wave sources. J Acoust Soc Am 2018;143(1):529. 10.1121/1.5021245.

[22] Acquaticci F, Yommi MM, Gwirc SN, Lew SE. Rapid Prototyping of Pyramidal Structured Absorbers for Ultrasound. Open Journal of Acoustics 2017;7:83–93. 10.4236/oja.2017.73008.

[23] Schoppe O, Pan C, Coronel J, Mai H, Rong Z, Todorov MI et al. Deep learning-enabled multi-organ segmentation in whole-body mouse scans. Nat Commun 2020;11(1). 10.1038/s41467-020-19449-7.

[24] IT’IS Foundation. Tissue Properties Database V4.1; 2022.

[25] Kroemer G, Galluzzi L, Kepp O, Zitvogel L. Immunogenic Cell Death in Cancer Therapy. Annu. Rev. Immunol. 2013;31(1):51–72. 10.1146/annurev-immunol-032712-100008.

[26] Ricci M, Dimitri M, Serio M, Corvi A. Morphological Analysis of US Treated PANC-1 Spheroids. Applied Sciences 2025;15(4):1707. 10.3390/app15041707.

[27] Uddin S, Komatsu D, Motyka T, Petterson S. Low-Intensity Continuous Ultrasound Therapies—A Systematic Review of Current State-of-the-Art and Future Perspectives. JCM 2021;10(12):2698. 10.3390/jcm10122698.

[28] Cohen G, Chandran P, Lorsung RM, Aydin O, Tomlinson LE, Rosenblatt RB et al. Pulsed-Focused Ultrasound Slows B16 Melanoma and 4T1 Breast Tumor Growth through Differential Tumor Microenvironmental Changes. Cancers 2021;13(7):1546. 10.3390/cancers13071546.

[29] Cohen G, Chandran P, Lorsung RM, Tomlinson LE, Sundby M, Burks SR et al. The Impact of Focused Ultrasound in Two Tumor Models: Temporal Alterations in the Natural History on Tumor Microenvironment and Immune Cell Response. Cancers 2020;12(2):350. 10.3390/cancers12020350.

[30] Xia Y, Yang M, Xiao X, Tang W, Deng J, Wu L et al. Low-intensity pulsed ultrasound activated the anti-tumor immunity by irradiating the spleen of mice in 4 T-1 breast cancer. Cancer Immunol Immunother 2024;73(3). 10.1007/s00262-023-03613-1.

[31] Aydin O, Chandran P, Lorsung RR, Cohen G, Burks SR, Frank JA. The Proteomic Effects of Pulsed Focused Ultrasound on Tumor Microenvironments of Murine Melanoma and Breast Cancer Models. Ultrasound in Medicine & Biology 2019;45(12):3232–45. 10.1016/j.ultrasmedbio.2019.08.014.

[32] Dai Z, Liu W-C, Chen X-Y, Wang X, Li J-L, Zhang X. Gasdermin D-mediated pyroptosis: mechanisms, diseases, and inhibitors. Front. Immunol. 2023;14. 10.3389/fimmu.2023.1178662.

[33] Fontana P, Du G, Zhang Y, Zhang H, Vora SM, Hu JJ et al. Small-molecule GSDMD agonism in tumors stimulates antitumor immunity without toxicity. Cell 2024;187(22):6165–6181.e22. 10.1016/j.cell.2024.08.007.

[34] Matsuhashi S, Manirujjaman M, Hamajima H, Ozaki I. Control Mechanisms of the Tumor Suppressor PDCD4: Expression and Functions. IJMS 2019;20(9):2304. 10.3390/ijms20092304.

[35] Sun J, Cui H, Gao Y, Pan Y, Zhou K, Huang J et al. TGF-α Overexpression in Breast Cancer Bone Metastasis and Primary Lesions and TGF-α Enhancement of Expression of Procancer Metastasis Cytokines in Bone Marrow Mesenchymal Stem Cells. BioMed Research International 2018;2018:1–10. 10.1155/2018/6565393.

[36] Shi W, Pan Y, Rathod B, Wang Y, Wang Z, Shen J et al. TGF-á/EGFR-mediated lymphatic metastasis reveals a repositionable therapeutic target in breast cancer. npj Breast Cancer 2026;12(1). 10.1038/s41523-026-00941-0.

[37] Ghiringhelli F, Apetoh L, Tesniere A, Aymeric L, Ma Y, Ortiz C et al. Activation of the NLRP3 inflammasome in dendritic cells induces IL-1beta-dependent adaptive immunity against tumors. Nat Med 2009;15(10):1170–8. 10.1038/nm.2028.

[38] Moossavi M, Parsamanesh N, Bahrami A, Atkin SL, Sahebkar A. Role of the NLRP3 inflammasome in cancer. Mol Cancer 2018;17(1). 10.1186/s12943-018-0900-3.

[39] Chen Y, Zhang J, Cui W, Silverstein RL. CD36, a signaling receptor and fatty acid transporter that regulates immune cell metabolism and fate. Journal of Experimental Medicine 2022;219(6). 10.1084/jem.20211314.

[40] Waight JD, Hu Q, Miller A, Liu S, Abrams SI. Tumor-Derived G-CSF Facilitates Neoplastic Growth through a Granulocytic Myeloid-Derived Suppressor Cell-Dependent Mechanism. PLoS ONE 2011;6(11):e27690. 10.1371/journal.pone.0027690.

[41] Gabrilovich DI, Nagaraj S. Myeloid-derived suppressor cells as regulators of the immune system. Nat Rev Immunol 2009;9(3):162–74. 10.1038/nri2506.

[42] Veglia F, Sanseviero E, Gabrilovich DI. Myeloid-derived suppressor cells in the era of increasing myeloid cell diversity. Nat Rev Immunol 2021;21(8):485–98. 10.1038/s41577-020-00490-y.

[43] Aslan V, Karabörk Kılıç AC, Özet A, Üner A, Günel N, Yazıcı O et al. The role of spleen volume change in predicting immunotherapy response in metastatic renal cell carcinoma. BMC Cancer 2023;23(1). 10.1186/s12885-023-11558-y.

[44] Galland L, Lecuelle J, Favier L, Fraisse C, Lagrange A, Kaderbhai C et al. Splenic Volume as a Surrogate Marker of Immune Checkpoint Inhibitor Efficacy in Metastatic Non Small Cell Lung Cancer. Cancers 2021;13(12):3020. 10.3390/cancers13123020.

[45] Castagnoli F, Doran S, Lunn J, Minchom A, O’Brien M, Popat S et al. Splenic volume as a predictor of treatment response in patients with non-small cell lung cancer receiving immunotherapy. PLoS ONE 2022;17(7):e0270950. 10.1371/journal.pone.0270950.

[46] Balkwill F. Tumour necrosis factor and cancer. Nat Rev Cancer 2009;9(5):361–71. 10.1038/nrc2628.

[47] Dinarello CA. Overview of the IL −1 family in innate inflammation and acquired immunity. Immunological Reviews 2018;281(1):8–27. 10.1111/imr.12621.

[48] Keerthi Raja MR, Gupta G, Atkinson G, Kathrein K, Armstrong A, Gower RM et al. Host-derived interleukin-1á drives tumor immunosuppression by reprogramming tumor-associated myeloid cells. npj Breast Cancer 2026;12(1):26. 10.1038/s41523-026-00890-8.

[49] Ouyang W, Rutz S, Crellin NK, Valdez PA, Hymowitz SG. Regulation and Functions of the IL-10 Family of Cytokines in Inflammation and Disease. Annu. Rev. Immunol. 2011;29(1):71–109. 10.1146/annurev-immunol-031210-101312.

