## Supplementary Information for "Simulation-guided non-thermal low-intensity ultrasound reprograms the tumor immune microenvironment and engages systemic antitumor immunity in a syngeneic orthotopic mouse model of breast cancer"

### Section S1. Supplementary experimental information

#### S1.1. *In vitro* ultrasound setup

The customized setup (**Fig. 1**) consisted of a water tank, a microplate holder, an acoustic absorber, a heater and a custom ultrasound device, that ensures repeatable experimental conditions and allows for fine control of the ultrasonic field and of the temperature, as outlined below. The polymethyl methacrylate (PMMA) water tank has a circular recess milled at the center of the base sheet to accommodate the ultrasonic transducer, reducing the wall thickness from 10 mm to 2 mm at that location. To ensure a consistent dish placement, a sample holder was 3D-printed with glycol-modified polyethylene terephthalate filament (Ultimaker PETG, Ultimaker B.V., Geldermalsen, The Netherlands) and registered with the tank walls. With the ultrasonic transducer positioned at the bottom, acoustic waves propagating vertically towards the water-air interface and reflections at that interface would cause a standing wave along the vertical axis. This was prevented by placing a pyramidal acoustic absorber between the water–air interface and the microplates. Its geometry was designed to minimize acoustic reflections based on previously described principles [1], and it was cast from Aptflex F36 (Precision Acoustics Ltd, Dorchester, United Kingdom) material following the manufacturer's casting instructions. For best results, a positive mold was 3D-printed on a Formlabs Form 3B with black resin (Formlabs, Somerville MA, United States of America). Stereolithography printing was chosen because fused-filament printers, such as the Ultimaker printers used for the petri dish holders mentioned above, lack the resolution required for this application. A negative mold was then created by casting ESSIL 125 silicone (Axson Technologies, Saint-Ouen-l'Aumône, France) into the 3D-printed mold. After curing, the Aptflex F36 compound was poured into the silicone mold. The water temperature was maintained with a heater (ProfiCook PC-SV 1126, Clatronic International GmbH, Kempen, Germany) clipped into a corner of the tank.

#### S1.2. Customized ultrasound-control system

##### Control electronics

A customized ultrasound system was developed to provide stable and reproducible continuous-wave excitation under the experimental conditions used in this study. The system comprised an enclosure containing the control electronics and a detachable 1MHz transducer. Device specifications and ultrasound exposure conditions were reported following published guidance for acoustic-output reporting [2]. The control electronics drove the transducer using a sinusoidal voltage of fixed frequency and controlled amplitude. The waveform-generation circuit comprised an AD9834 direct digital synthesizer and an AD8130 differential amplifier (Analog Devices, Wilmington, MA, USA). The resulting signal was further amplified using an ADA4870 linear power stage (Analog Devices) with sufficient bandwidth and local feedback to drive the transducer with low distortion. A mean-responding power detector (AD8361, Analog Devices) monitored a current-related signal from the transducer. A microcontroller-based feedback algorithm used this measurement to adjust the amplitude of the sinusoidal drive voltage and stabilize the electrical power delivered to the transducer. The system continuously monitored its operating state and generated a warning or deactivated ultrasound emission when operation deviated from the predefined conditions.

##### Transducer and coupling interface

Ultrasound was generated using a disk-shaped piezoelectric transducer with a nominal resonance frequency of 1MHz (SMMSG25F1000, STEMiNC, Miami, FL, USA). The active element was composed of modified PZT-4 material and had a diameter of 25 mm. The transducer was mounted in a custom housing designed for attachment to the underside of the water tank. Its coupling interface consisted of a flat, disk-shaped stainless-steel surface that provided mechanical coupling between the transducer and the tank floor. For all experiments reported in this study, the system was operated at 1MHz in continuous-wave mode (100% duty cycle). The electrical drive-power set point was 4.910 W. This value represents the electrical power delivered by the control electronics to the transducer and should not be interpreted as the acoustic output power or acoustic intensity at the sample plane. The system was configured and monitored using a dedicated computer application communicating with the control electronics through a USB connection. The application enabled selection of the operating frequency and exposure mode, adjustment of the electrical drive-power set point, and continuous monitoring of device operation throughout ultrasound exposure.

**Relative output-linearity assessment**

The relative linearity and stability of the ultrasound output were evaluated using an ad hoc radiation-force-balance setup based on IEC 61161:2013, “Ultrasonics—Power measurement—Radiation force balances and performance requirements”. The radiation-force-balance assembly was mounted on the experimental water tank and used to assess changes in effectively coupled acoustic output as a function of the electrical drive-power set point. Because the acoustic absorber available for this assessment had not been independently calibrated or certified for absolute radiation-force-balance measurements, the setup was not used to determine absolute acoustic power. It was used exclusively to assess the relative linearity and stability of the coupled ultrasound output. For the 1MHz transducer, the measured response was approximately linear at electrical drive-power settings of up to 5 W. Saturation became apparent at settings between 5 and 8 W. The electrical drive-power set point used in the present study, 4.910 W, therefore remained within the experimentally determined linear operating range.

#### S1.3. Analysis of RNA sequencing data

Sequencing data were analyzed using Qiagen CLC Genomics Server 23.0.5 (Qiagen, Hilden, Germany) with the following workflow. Raw reads were trimmed to remove adapter sequences and low-quality bases, allowing a maximum of two ambiguous nucleotides. Reads were initially mapped to the SILVA 132 rRNA database to assess rRNA content, then aligned to the mouse genome GRCm38 (ENSEMBL v98). For unsupervised analyses (**Fig. 9A, B**), genes were considered "expressed" if they had at least 10 reads in one group of replicates, and a variance stabilizing transformation (vst) was applied using the vst function from DESeq2 v1.28.1 [3] with blind=TRUE to calculate variance independently of pre-defined groups. For the pairwise comparison between untreated and LIU-treated groups, differentially expressed genes (DEGs) were identified using DESeq2's Wald test, with multiple testing corrected via the Benjamini–Hochberg algorithm to control the false discovery rate (FDR). Genes with $p < 0.05$ and FDR < 0.05 were considered statistically significant and used for further functional analysis. To identify enriched biological processes and gene ontologies (GOs), up- and down-regulated DEGs were analyzed using Enrichr [4]. ClueGO (v2.5.8) [5] in Cytoscape (v3.9.1) [6] was used for biological network enrichment and Ingenuity Pathway Analysis (IPA) was employed to predict significantly affected canonical pathways and upstream regulators.

### References

1. Acquaticci F, Yommi MM, Gwirc SN, Lew SE. Rapid Prototyping of Pyramidal Structured Absorbers for Ultrasound. Open Journal of Acoustics. 2017;7:83–93. doi:10.4236/oja.2017.73008.
2. Edmonds PD, Abramowicz JS, Carson PL, Carstensen EL, Sandstrom KL. Guidelines for Journal of Ultrasound in Medicine Authors and Reviewers on Measurement and Reporting of Acoustic Output and Exposure. J of Ultrasound Medicine. 2005;24:1171–9. doi:10.7863/jum.2005.24.9.1171.
3. Love MI, Huber W, Anders S. Moderated estimation of fold change and dispersion for RNA-seq data with DESeq2. Genome Biol. 2014;15:550. doi:10.1186/s13059-014-0550-8.
4. Chen EY, Tan CM, Kou Y, Duan Q, Wang Z, Meirelles GV, et al. Enrichr: interactive and collaborative HTML5 gene list enrichment analysis tool. BMC Bioinformatics. 2013;14:128. doi:10.1186/1471-2105-14-128.
5. Bindea G, Mlecnik B, Hackl H, Charoentong P, Tosolini M, Kirilovsky A, et al. ClueGO: a Cytoscape plug-in to decipher functionally grouped gene ontology and pathway annotation networks. Bioinformatics. 2009;25:1091–3. doi:10.1093/bioinformatics/btp101.
6. Shannon P, Markiel A, Ozier O, Baliga NS, Wang JT, Ramage D, et al. Cytoscape: a software environment for integrated models of biomolecular interaction networks. Genome Res. 2003;13:2498–504. doi:10.1101/gr.1239303.

**Section S2. Supplementary Figures**

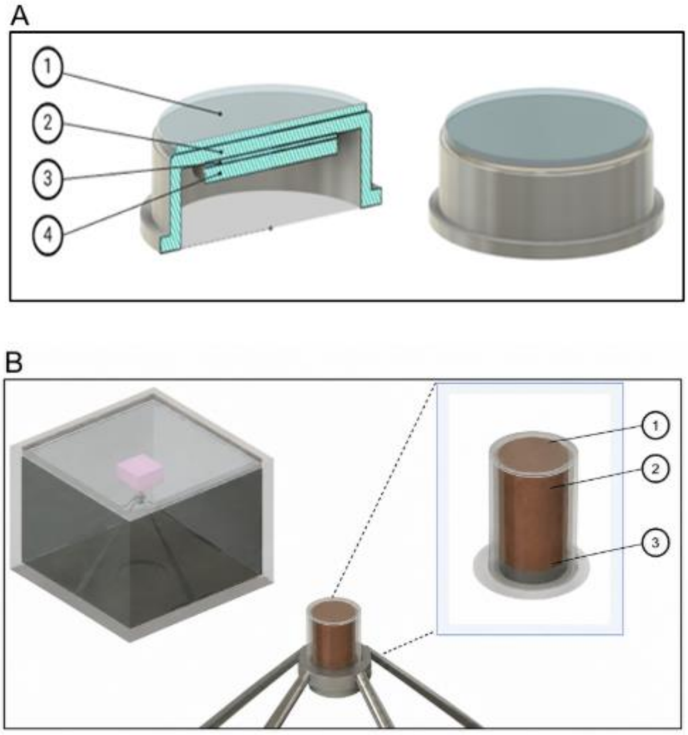

**Fig. S1.** Computational modelling of in vitro conditions. (A) Simulated layers of the transducer. 1: coupling gel, 2: steel housing, 3: glue, 4: piezo element. (B) Left: 96 well setup in the water tank (PMMA) with an absorber (pink). Middle: microplate held in position by the sample holder. Right: 1: Parafilm, 2: microplate, 3: cell culture layer.

**A**

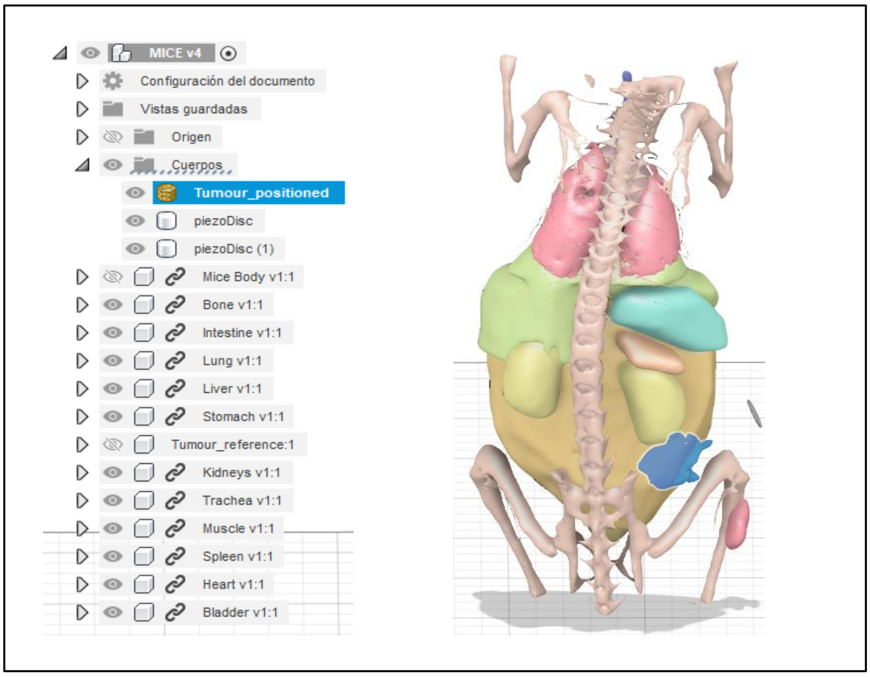

**B**

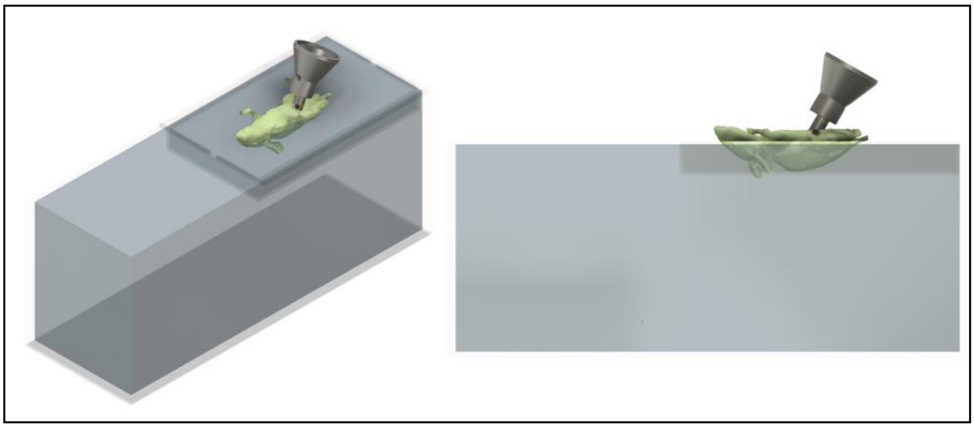

**Fig. S2.** Computational modelling of in vivo conditions. (A) Interior of mouse showing the body parts obtained from the segmentation: bones, intestine, lungs, liver, stomach, kidneys, trachea, muscle, spleen, heart, bladder. (B) simulated configuration including water tank, mouse and transducer.

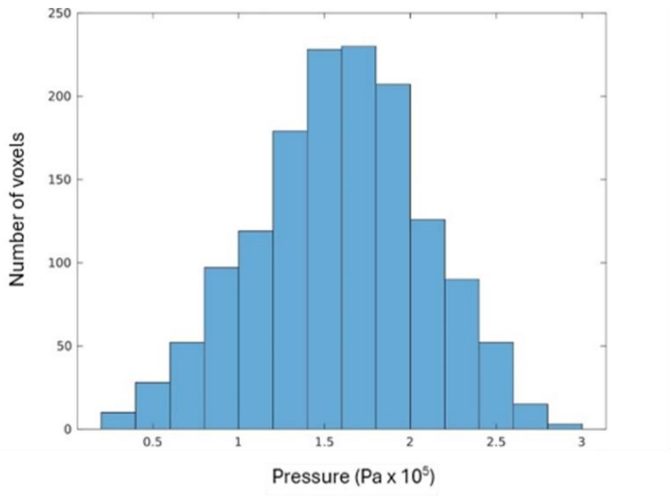

**Fig. S3.** Distribution of simulated acoustic-pressure amplitudes across the cell-culture plane. Histogram showing the acoustic-pressure amplitudes calculated for individual voxels within the simulated cell layer at the optimized transducer-to-sample distance of 94mm. Simulations were performed at 1MHz, an input intensity of 1W/cm², and a 100% duty cycle and were continued until multiple boundary reflections produced a stable interference pattern. The distribution of voxel-level pressure amplitudes supports a relatively homogeneous acoustic exposure across the cell-culture plane without a distinct population of localized high-pressure voxels.

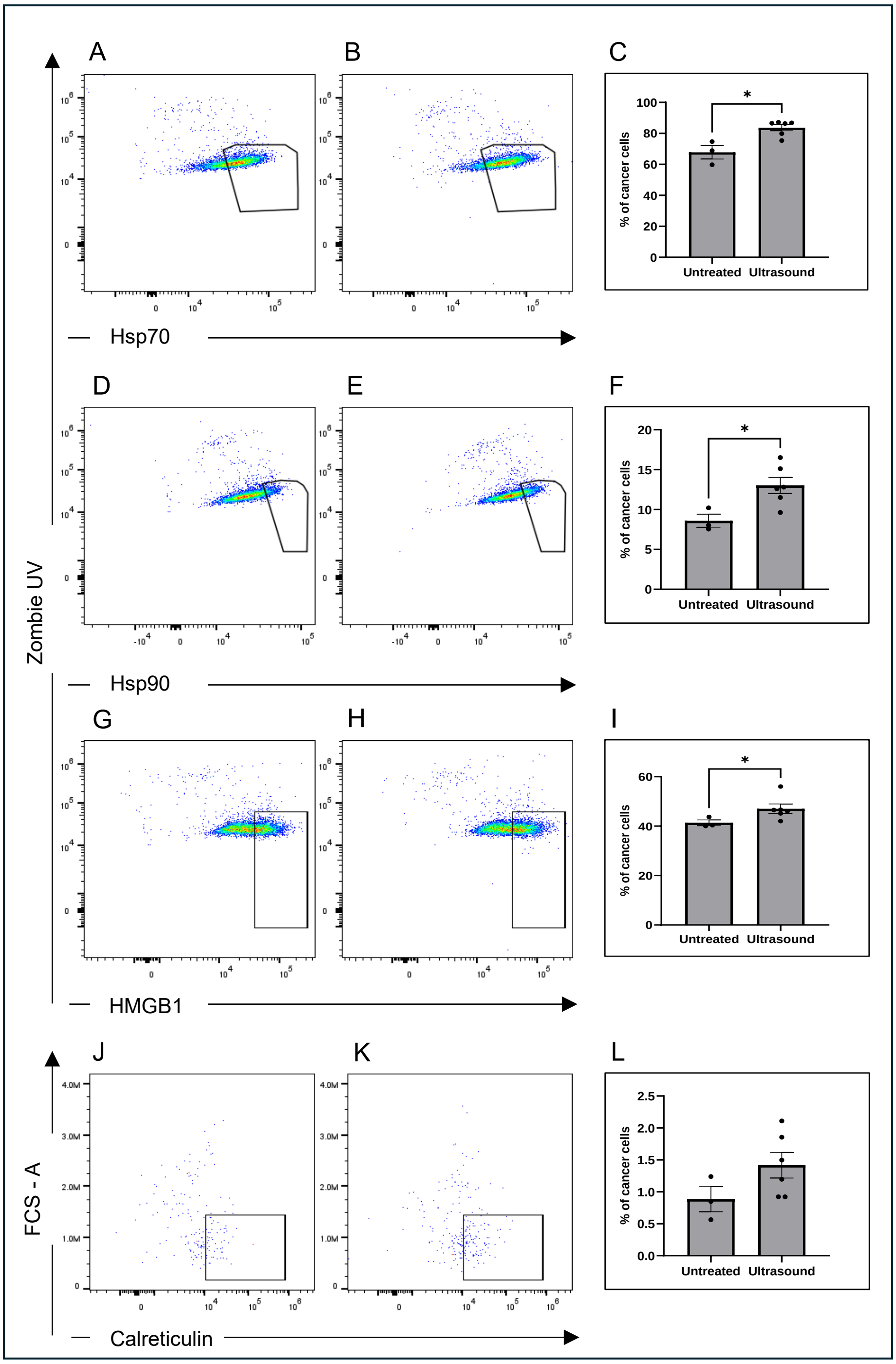

**Figure S4.** Flow cytometry analysis of low-intensity ultrasound (LIU)-mediated effects on 4T07 spheroids.

4T07 spheroids were treated at 1MHz, 1W/cm², 100% duty cycle, 20min. Pseudocolor plots display the distribution of Hsp70 (A, B), Hsp90 (D, E), HMGB1 (G, H), and calreticulin (J, K) in untreated and ultrasound-treated spheroids. Bar graphs (C, F, I, L) show the percentage of cells positive for each marker. Results are presented as mean ± SEM, with individual data points shown as black dots. Statistical analysis was performed using Welch's t-test for Hsp90 and calreticulin, while the nonparametric Mann-Whitney test was used for Hsp70 and HMGB1. Statistical significance is indicated by asterisks (*$p < 0.05$).

### Section S3. Supplementary Tables

**Table S1.** Gene-level references for transcriptomic findings

| Gene(s) | Mouse symbol | Biological role in context | Reference(s) |
|---|---|---|---|
| *Mmp1a, Mmp10* | Matrix metalloproteinases | ECM degradation; prerequisite for local invasion and metastasis | Justilien et al., 2012; Gabasa et al., 2021 |
| *Tgfa* | Transforming growth factor alpha | Tumor-stroma signaling; immune evasion axis | Jhappan et al., 1990; Justilien et al., 2012; Niu et al., 2022 |
| *Pdcd4* | Programmed cell death 4 | Tumor suppressor; inhibits translation of pro-survival factors | Matsuhashi et al., 2019 |
| *Casp4* | Caspase 4 | Non-canonical inflammasome; cleavage of Gasdermin D triggering pyroptosis | Dai et al., 2023; Li et al., 2024 |
| *Gsdmd* | Gasdermin D | Pyroptosis effector; pore formation and inflammatory cell death | Yu et al., 2021; Dai et al., 2023 |
| *Nlrp3* | NOD-like receptor protein 3 | Inflammasome sensor; promotes dendritic cell maturation and CD8⁺ T cell activation | Hanamsagar et al., 2011; Kurdi et al., 2018 |
| *Cd36* | Cluster of differentiation 36 | Scavenger receptor; ROS generation and pro-inflammatory cytokine production | Chen et al., 2022; Preedy et al., 2024 |
| *Tlr2, Nod1, Nod2* | Pattern-recognition receptors | Innate immune sensing; antigen presentation, neutrophil and macrophage recruitment | Feng et al., 2019; Geijtenbeek & Gringhuis, 2009; Omaru et al., 2022 |
| *Clec4a1, Clec9a* | C-type lectin receptors | Sensing dying cells; cross-presentation of tumor antigens to CD8⁺ T cells | Schreibelt et al., 2012; Uto et al., 2016; Kircheis & Planz, 2023 |
| *Cd5l* | CD5 antigen-like | Anti-inflammatory macrophage marker; immune regulation | Menon et al., 2023 |
| *Cxcl5* | C-X-C motif chemokine ligand 5 | Neutrophil recruitment | Takimoto-Sato et al., 2023; Zhang et al., 2024 |
| TLR signaling (general) | *Tlr1, Tlr7–Tlr9, Tlr11–Tlr13* | Activation of dendritic cells, macrophages, T lymphocytes; antigen presentation | Duan et al., 2022 |
| *Cd3d, Cd3e, Cd3g* | T cell receptor complex | T cell activation, infiltration, cytotoxic function | Cibrián & Sánchez-Madrid, 2017; Waldman et al., 2020 |
| *Cd4, Cd8a, Cd8b1* | T cell co-receptors | Helper and cytotoxic T cell identity and function | Wang et al., 2012; Ou et al., 2023; Koh et al., 2023 |
| *Cd69, Il2ra* | Early T cell activation markers | T cell activation and proliferation | Hellstrom et al., 2001 |
| *Tnfrsf4* (OX40) | OX40 costimulatory receptor | T cell proliferation and survival | Croft et al., 2009 |
| *Icos* | Inducible T cell co-stimulator | T cell effector function and memory | Wikenheiser & Stumhofer, 2016 |
| *Ctla4* | Cytotoxic T-lymphocyte antigen 4 | Counter-regulatory checkpoint; T cell inhibition | Raskov et al., 2021 |
| *Cd19, Ms4a1* (CD20) | B cell surface markers | B cell identity and activation | Pavlasova & Mraz, 2020; Texido et al., 2000 |
| *Cd27, Tnfrsf13c* | B cell co-stimulatory receptors | B cell survival and antibody production | Smulski & Eibel, 2018 |
| *Blk* | B lymphocyte kinase | B cell receptor signaling | Grimsholm, 2023 |
| *Cxcl13* | B cell-attracting chemokine 1 | B cell recruitment; tertiary lymphoid structure formation | Zhao et al., 2024 |
| *Cxcl9, Cxcl11* | Interferon-inducible chemokines | Recruitment of T cells and NK cells | Tokunaga et al., 2018; Immler et al., 2024 |
| *Ccl21a, Ccr3* | CC chemokines | Recruitment of dendritic cells and T cells | Ghaffari & Rezaei, 2023; Yan et al., 2021 |

**Section S4. Supplementary Files**

**File S1.** List of the genes with up and down regulations, gene ontology (GO), Cytoscape analysis, and Ingenuity Pathway Analysis.
